# Calling differential DNA methylation at cell-type resolution: addressing misconceptions and best practices

**DOI:** 10.1101/2021.02.14.431168

**Authors:** Elior Rahmani, Brandon Jew, Regev Schweiger, Brooke Rhead, Lindsey A. Criswell, Lisa F. Barcellos, Eleazar Eskin, Saharon Rosset, Sriram Sankararaman, Eran Halperin

## Abstract

We benchmarked two approaches for the detection of cell-type-specific differential DNA methylation: Tensor Composition Analysis (TCA) and a regression model with interaction terms (CellDMC). Our experiments alongside rigorous mathematical explanations show that TCA is superior over CellDMC, thus resolving recent criticisms suggested by Jing et al. Following misconceptions by Jing and colleagues with modelling cell-type-specificity and the application of TCA, we further discuss best practices for performing association studies at cell-type resolution. The scripts for reproducing all of our results and figures are publicly available at github.com/cozygene/CellTypeSpecificMethylationAnalysis.

---

Calling differential DNA methylation at a cell-type level from tissue-level bulk data has recently become a question of interest^1–6^, and thus far, two main different approaches have been suggested for the task. The first approach employs a standard regression analysis while including and evaluating interaction terms (i.e. multiplicative terms) between cell-type proportions and the phenotype of interest. This approach has long been suggested and repeatedly established in the context of cell-type-specific differential expression analysis^7,8^, and it was recently proposed in the context of methylation by Zheng et al. (a method called CellDMC)^2^; notably, this approach has been independently applied to methylation by other groups as well^5,6^. We recently presented a second approach, called Tensor Composition Analysis (TCA). The TCA framework is based on a novel method we developed and applied for modelling cell-type-specific variability; particularly, we presented it in the context of detecting differential methylation at cell-type-specific resolution^3^. In response to the original paper introducing TCA, Jing et al.^9^ made multiple claims about the utility of TCA and its performance compared with CellDMC. We argue that these claims stem from methodological misconceptions and improper application of TCA. We hereby address these claims and clarify the distinction between TCA and CellDMC, with the hope that this will resolve confusions about the TCA framework and will help users to apply it correctly.

First and foremost, we stress out that in principle TCA cannot be inferior to CellDMC when compared objectively: TCA is a more expressive model than CellDMC, as it makes more general assumptions about the variation of cell-type-specific methylation. Therefore, for large enough data (at least 60 samples as we later show), TCA is expected to be the better choice in general. To illustrate the difference between the methods, consider a case/control study design. In that case, CellDMC assumes a fixed effect between cases and controls as the only variation at the cell-type level; this corresponds to the unrealistic assumption that all individuals within a group (i.e. cases or controls) have the exact same methylome. TCA improves upon this by modeling the variation of cell-type-specific methylation across individuals (i.e. even within the same group). We show this theoretically by revealing the mathematical relation between TCA and CellDMC, which yields CellDMC as a degenerate case of the more general TCA model (Supplementary Note). In order to demonstrate this result empirically, we conducted a simulation study while following the same simulation setup and evaluation metrics outlined by Jing and colleagues^9^. As we show next, we observe that TCA improves upon CellDMC under all scenarios proposed by Jing et al. Yet, due to two key misconceptions we detail below, Jing et al. did not provide an objective comparison of TCA and CellDMC, which led to a major discrepancy in results.

The first misconception derives from an inconsistent selection of statistical tests for comparison. Jing et al. evaluated TCA and CellDMC using two conceptually different types of statistical tests. While CellDMC fits a *marginal conditional* model, wherein the effects of all cell types are estimated jointly and then tested in each cell type, in their application of TCA they only considered a *marginal* model, wherein the effect of each cell type is estimated and tested marginally (i.e. separately), irrespective of other cell types. This distinction created a bias in their reporting, as these two types of tests inherently lead to different results; this property, which is not unique to TCA, essentially reduces to confusions about basic concepts in regression (Supplementary Note). Admittedly, we acknowledge that we should have provided a thorough discussion about the differences between *marginal conditional* and *marginal* tests in the TCA paper; of note, the software we provided with the publication clarified the differences.

The second misconception relates to inconsistency with the underlying biological assumptions taken in the application of TCA and CellDMC. Particularly, Jing et al. mixed up two distinct biological concepts: while they followed in CellDMC and their simulation study the assumption that methylation levels are *affected* by the phenotype of interest (or by a mediating component thereof; denote this assumption by X|Y), at the same time, they applied TCA in a way that corresponds to the assumption that methylation levels *affect* the phenotype of interest (or a mediating component thereof; denote this assumption by Y|X). Indeed, our original demonstration of TCA focused on the Y|X assumption, however, the TCA framework can also properly accommodate the X|Y assumption (see “Applying TCA to epigenetic association studies” in the Methods section of our original TCA paper^3^; Supplementary Note). For a comprehensive list of the statistical tests and biological assumptions implemented in TCA and in CellDMC see Supplementary Table 1.

Evaluating both methods under the same biological model and statistical test used in CellDMC (i.e. marginal conditional tests under the assumption X|Y) renders TCA as the (mildly) better performing method under all four scenarios and all three evaluation metrics that were considered by Jing et al. (Figure 1), as expected per our theoretical result (Supplementary Note). Repeating this analysis using different sample sizes shows a slight advantage for CellDMC over TCA in data with less than 60 individuals (Supplementary Figures 3 and 4). Further simulating phenotypes to be statistically affected by methylation (i.e. setting Y|X as the true model, rather than X|Y as in Jing et al.) results in a substantial decrease in specificity and precision for CellDMC (with the benefit of a mild increase in sensitivity) compared to TCA (Figure 2). This can be explained by the fact that TCA can properly accommodate the assumption Y|X, the true biological model in this case, whereas CellDMC is bound to assume X|Y. Notably, applying TCA under the wrong biological assumption in this case (i.e. assuming X|Y) performs better than CellDMC, reflecting better robustness of TCA to model misspecification (Supplementary Figure 1).

**Figure 1:**
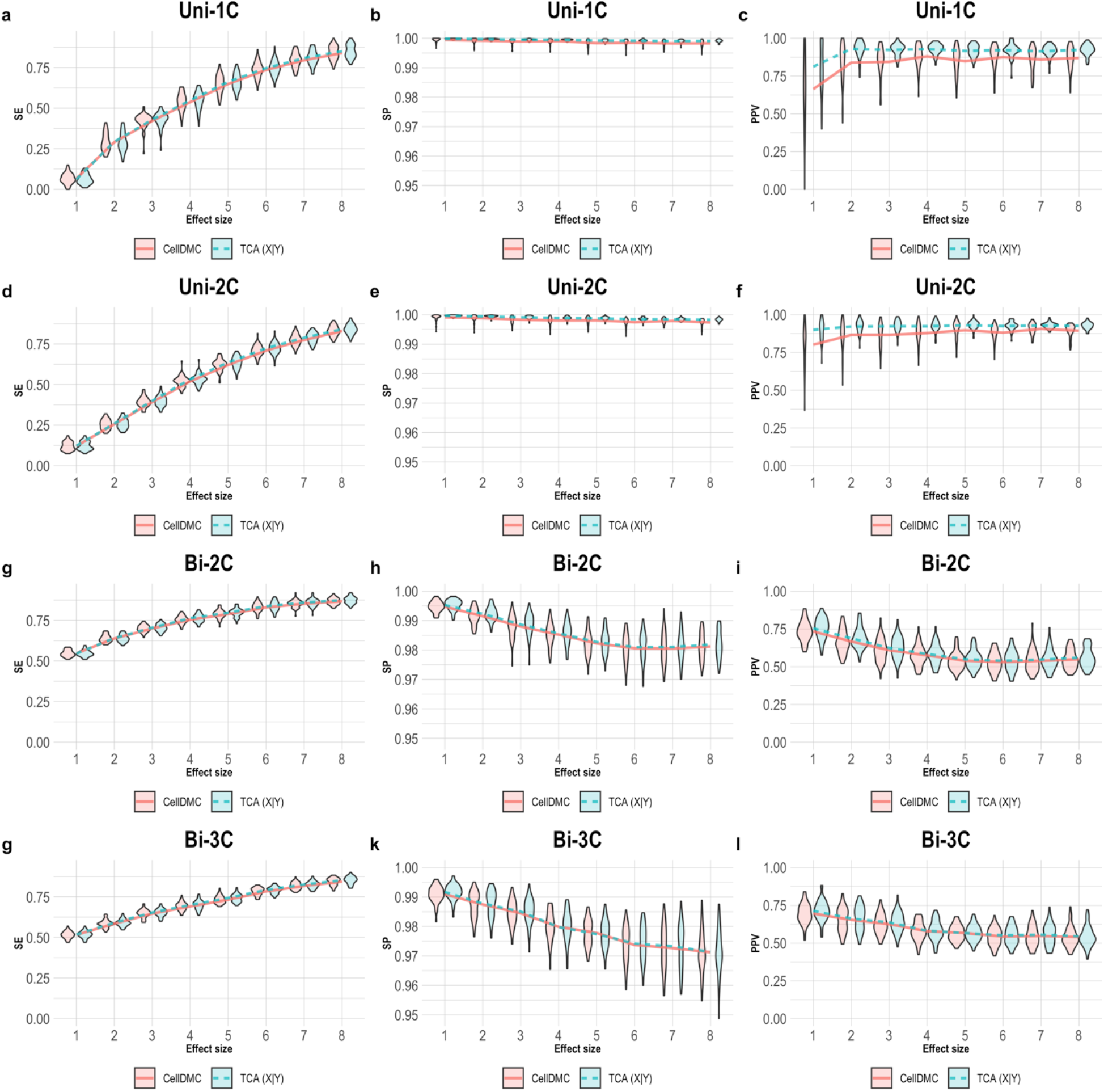
Evaluation of TCA and CellDMC in the case where the phenotype *affects* methylation (X|Y). (a)-(c) Comparison of the sensitivity (SE), specificity (SP), and precision (positive predictive value; PPV) to detect differentially methylated cell-types as a function of the association effect size, under the scenario where a single cell type out of 6 blood cell types is altered in cases versus controls (Uni-1C). (d)-(f) as in Uni-1C, only for the scenario where two cell types are altered in the same direction (Uni-2C). (g)-(i) as in Uni-2C, only for the scenario where the cell types are altered in opposite directions (Bi-2C). (j)-(l) as in Bi-2C, only for three cell types (Bi-3C). Results are shown across 50 simulated datasets using violin plots; solid lines represent median values. TCA was executed under the assumption X|Y (TCA X|Y).

**Figure 2:**
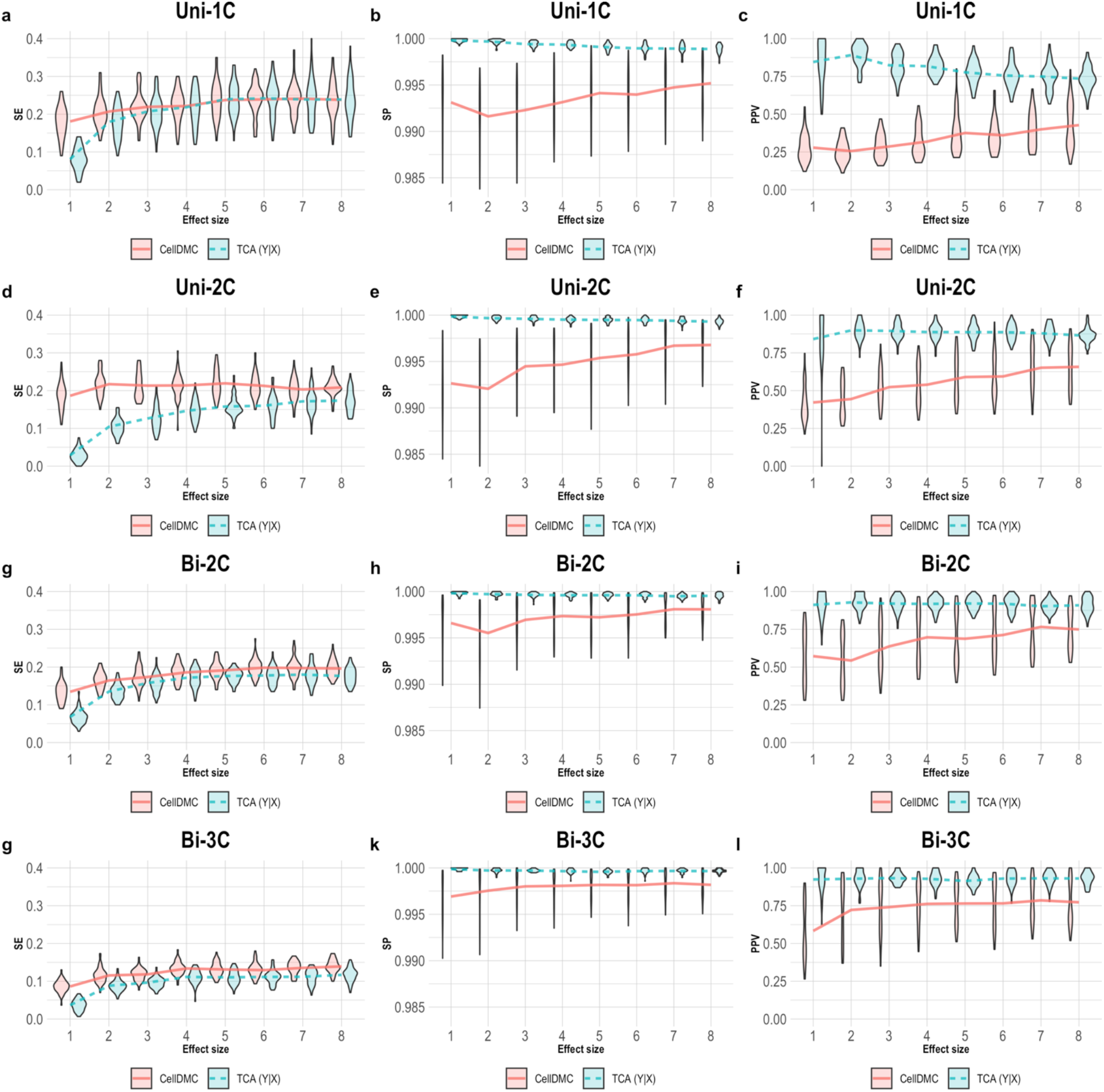
Evaluation of TCA and CellDMC in the case where the phenotype is *affected* by methylation (Y|X). (a)-(c) Comparison of the sensitivity (SE), specificity (SP), and precision (positive predictive value; PPV) to detect differentially methylated cell-types as a function of the association effect size, under the scenario where a single cell type out of 6 blood cell types is altered in cases versus controls (Uni-1C). (d)-(f) as in Uni-1C, only for the scenario where two cell types are altered in the same direction (Uni-2C). (g)-(i) as in Uni-2C, only for the scenario where the cell types are altered in opposite directions (Bi-2C). (j)-(l) as in Bi-2C, only for three cell types (Bi-3C). Results are shown across 50 simulated datasets using violin plots; solid lines represent median values. TCA was executed under the assumption Y|X (TCA Y|X).

Jing et al. further report poor performance when applying TCA for estimating cell-type proportions in two whole-blood datasets. However, we could not reconstruct their results; in fact, we found that the TCA estimates are almost in par with reference-based estimates (Supplementary Figure 2). Regardless, TCA, which requires cell-type proportions as an input, should not always be set to re-estimate cell-type proportions. Specifically, the iterative procedure in TCA for re-estimating proportions was designed for scenarios where the input estimates are expected to be limited in quality (e.g., due to lack of appropriate methylation reference); our previous experiments verified the performance of TCA under such cases^3^.

Another criticism made by Jing et al. concerns the computational efficiency of TCA. We note that speeding up TCA was out of scope in the original paper, where we chose to focus on the development of the first-of-its-kind model-based deconvolution method that allows learning 3-dimensional signals from 2-dimensional data. However, speed is not an inherent limitation of TCA. In order to show this, we improved the runtime of TCA by an order of magnitude by implementing a simple and substantially faster optimization (Supplementary Note), resulting in runtime that is more comparable to CellDMC’s (Supplementary Figures 5,6).

Lastly, Jing et al. claim that TCA was not properly evaluated on real data. Particularly, they dismissed our evaluation of TCA using previously published data with rheumatoid arthritis (RA)^10^, claiming that RA has no reliable known associations that can be used as ground truth. This claim is unclear to us, given that most of the authors in Jing et al. suggested their analysis of the very same dataset as evidence for the utility of CellDMC^2^. Moreover, we indeed provided replication analysis for the multiple associations we found with TCA using independent RA data collected from sorted cells; notably, none of the associations that were reported by CellDMC in this experiment could be replicated in the sorted methylation data.

Jing and colleagues further present results with smoking status in two whole-blood datasets^10,11^, where they used previously reported cell-type-specific associations in 7 CpGs^12^ as ground truth for evaluation. However, as in their simulations, they did not provide a comparison of TCA with CellDMC using the same statistical test and under the same biological assumption, which would have revealed essentially the same results for TCA and CellDMC in this case (Figure 3a-b). Since any evaluation of sensitivity to detect true positives should be complemented with an evaluation of specificity, we further evaluated the performance of the two methods on the entire data (i.e. rather than just on the 7 CpGs). Our results show that while TCA is well calibrated at the epigenome-wide level, CellDMC tends to suffer from a severe inflation in test statistic, which indicates low specificity and precision (Figure 3d).

**Figure 3:**
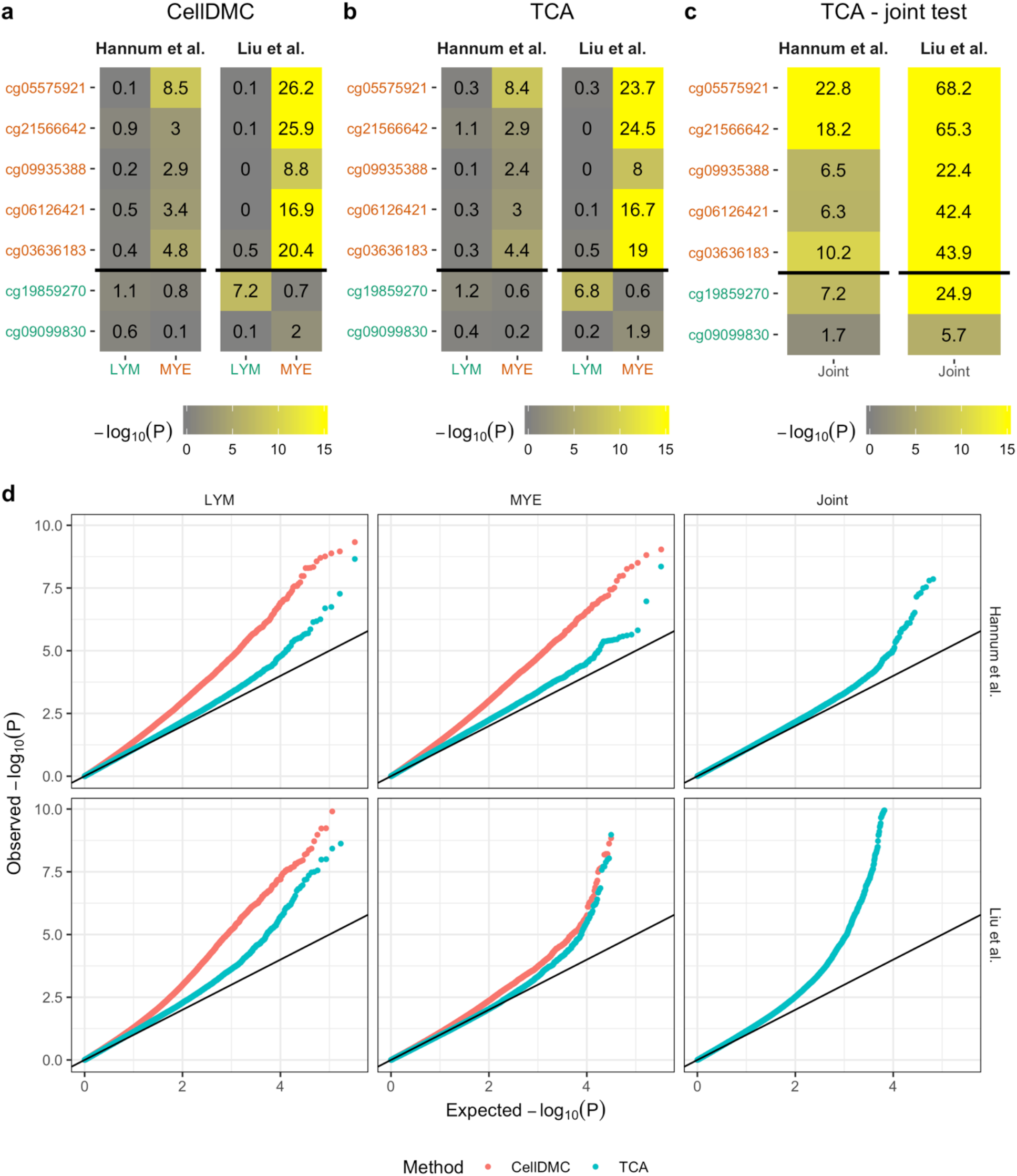
Evaluation of TCA and CellDMC in two independent whole-blood datasets with smoking. (a-c) Association tests were performed for each of 7 CpGs that were previously reported by Su et al. as exhibiting either myeloid-specific (in red) or lymphoid-specific (in green) associations with smoking status^12^. Results are displayed as heatmaps of the (negative-log transformed) p-values of the associations with myeloid cells (neutrophils and monocytes) and with lymphoid cells (T-cells, B-cells, and NK-cells) using (a) CellDMC, (b) TCA under the assumption X|Y (using the ‘tca’ function), and (c) TCA under the assumption X|Y, while using a joint test for tissue-level significance (using the ‘tca’ function). The latter achieves genome-wide significance (i.e. >6.98, assuming all 450K methylation array sites) in all but one CpG; calling the cell types that drive these associations using the results in (b) as a post-hoc analysis reveals the high-power of combining these two tests. (d) Results of an epigenome-wide analysis presented by quantile-quantile plots of the (negative-log transformed) p-values for the association tests in (a)-(c). Significant global deviation from the y=x line indicates an inflation arising from a badly specified model. Axes were truncated for visual purposes.

Of note, 7 out of the 14 tested CpGs across the two datasets with smoking did not achieve genome-wide significance, which would not have allowed de-novo detection of these associations in practice, presumably due to insufficient power. In order to address this, it is important to first appreciate that modeling the cell-type-specific nature of methylation is expected to benefit more types of analyses beyond calling for differentially methylated cell types. Particularly, compared to a standard regression analysis, TCA improves the detection of tissue-level associated CpGs via *joint* tests, wherein the effects of all cell types are tested jointly for their combined effect^3^. Such tissue-level tests can enable a powerful two-step approach of first detecting tissue-level associations followed by a post-hoc analysis of the associated CpGs at the cell-type level. Indeed, combining a tissue-level test for each CpG with a cell-type level post-hoc analysis, as allowed by TCA, correctly detects 10 out of the 14 smoking associated CpGs and cell types at a genome-wide significance level (Figure 3c). This shows that the detection of tissue-level associations is of primary interest, and methods such as TCA and CellDMC should not be evaluated solely on their ability to directly capture differentially methylated cell types as argued by Jing et al.

While we resolve their main concerns, we would like to commend Jing and colleagues for pursuing clarity and a continuous evaluation of methodologies, which is a key for developing best practices in research. Particularly, we acknowledge that precision of methods, which was not discussed in detail either in the TCA paper or in the CellDMC paper, should explicitly be considered as an evaluation metric whenever possible. Further, we agree that the reporting of results should always strive for clarity. Yet, in simulation studies that involve a great amount of detail, reproducibility is often challenging in practice. For that reason, here, we take a step forward and release the code for reconstructing our entire analysis, and we encourage others in the community to consider this approach in their publications to improve reproducibility.

In summary, we provide both empirical and theoretical evidence that for large enough sample sizes (at least 60), TCA is superior over CellDMC when it is applied under the assumptions taken in CellDMC, with the additional benefit of allowing to accommodate and therefore better handle different assumptions that are not allowed by CellDMC. Following these results and our demonstration that computational efficiency is not a limitation of TCA, we recommend that TCA should always be preferred over CellDMC, as long as the sample size is above 60.

Finally, as a practical note, performing marginal tests, as in Jing et al.’s application of TCA, substantially improves power over the alternatives, however, at the cost of a considerable decrease in precision (Supplementary Figures 7, 8, and 9). We therefore reiterate our previous recommendation to complement large data generation with small samples of sorted methylation data^3^. Such data can address the low precision limitation of the highly powerful marginal tests by providing a way to experimentally replicate associations at a cell-type-specific resolution. In the absence of such data for validation, it is advised to use the less powerful yet more precise alternative tests provided in TCA (see Supplementary Note for a comprehensive discussion and recommendations).

## Notes

https://github.com/cozygene/CellTypeSpecificMethylationAnalysis

### Competing Interest Statement

The authors have declared no competing interest.

