## Supplementary Information for "Calling differential DNA methylation at cell-type resolution: addressing misconceptions and best practices"

### 1 Supplementary Tables

| Model direction | Model | Statistical test | TCA implementation | CellDMC implementation |
|---|---|---|---|---|
| $Y\|X$ | $y_i = \sum_{h=1}^{k} Z^i_{hj}\beta_{hj} + e_i$<br>$e_i \sim N(0, \phi^2)$ | *marginal conditional*<br>for each cell type $h$:<br>$H_0 : \beta_{hj} = 0$<br>$H_1 : \beta_{hj} \neq 0$ | function `tcareg`<br>with `test='marginal_conditional'` | NA |
| | $y_i = Z^i_{hj}\beta_{hj} + e_i$<br>$e_i \sim N(0, \phi^2)$ | *marginal*<br>$H_0 : \beta_{hj} = 0$<br>$H_1 : \beta_{hj} \neq 0$ | function `tcareg`<br>with `test='marginal'` | NA |
| | $y_i = \sum_{h=1}^{k} Z^i_{hj}\beta_{hj} + e_i$<br>$e_i \sim N(0, \phi^2)$ | *joint*<br>$H_0 : \forall h, \beta_{hj} = 0$<br>$H_1 : \exists h, \beta_{hj} \neq 0$ | function `tcareg`<br>with `test='joint'` | NA |
| | $y_i = \sum_{h \in S_1} Z^i_{hj}\beta_{hj} + e_i$<br>$e_i \sim N(0, \phi^2)$ | *custom*<br>for $S_0 \subset S_1 \subseteq \{1, ..., k\}$:<br>$H_0 : \forall h \in S_1 \setminus S_0, \beta_{hj} = 0$<br>$H_1 : \exists h \in S_1 \setminus S_0, \beta_{hj} \neq 0$ | function `tcareg`<br>with `test='custom'` | NA |
| | $y_i = \sum_{h=1}^{k} Z^i_{hj}\beta_j + e_i$<br>$e_i \sim N(0, \phi^2)$ | *single effect size*<br>$H_0 : \beta_j = 0$<br>$H_1 : \beta_j \neq 0$ | function `tcareg`<br>with `test='single_effect'` | NA |
| $X\|Y$ | $\forall h : Z^i_{hj} = \mu_{hj} + y_i\gamma^j_h + \epsilon^i_{hj}$<br>$\epsilon^i_{hj} \sim N(0, \sigma^2_{hj})$ | *marginal conditional*<br>for each cell type $h$:<br>$H_0 : \gamma^j_h = 0$<br>$H_1 : \gamma^j_h \neq 0$ | function `tca` | function `CellDMC` * |
| | $\forall h : Z^i_{hj} = \mu_{hj} + y_i\gamma^j_h + \epsilon^i_{hj}$<br>$\epsilon^i_{hj} \sim N(0, \sigma^2_{hj})$ | *joint*<br>$H_0 : \forall h, \gamma^j_h = 0$<br>$H_1 : \exists h, \gamma^j_h \neq 0$ | function `tca` | NA ** |

Table 1: A summary of the statistical tests available in the TCA R package (version 1.2.0 and above; available on CRAN) and in CellDMC (part of the EpiDISH R package; available on Bioconductor). Notations: we denote the assumption that methylation *affects* the phenotype by $Y|X$ and the assumption that methylation is *affected* by the phenotype by $X|Y$, $Z^i_{hj}$ as the cell-type-specific methylation of individual $i$ at methylation site $j$ in cell type $h$, and $y_i$ as the value of the phenotype under test in individual $i$. Covariates are ignored in all models for simplicity. *Note that CellDMC does not explicitly model the cell-type-specific methylation $Z^i_{hj}$; it rather implicitly tries to capture $Z^i_{hj}$ by considering interaction terms between the cell-type proportions and the phenotype under test. ** A joint test was not implemented for CellDMC, however, the CellDMC model can in principle be modified to employ a partial F-test on the coefficients of the interaction terms under test.

#### 2 Supplementary Figuers

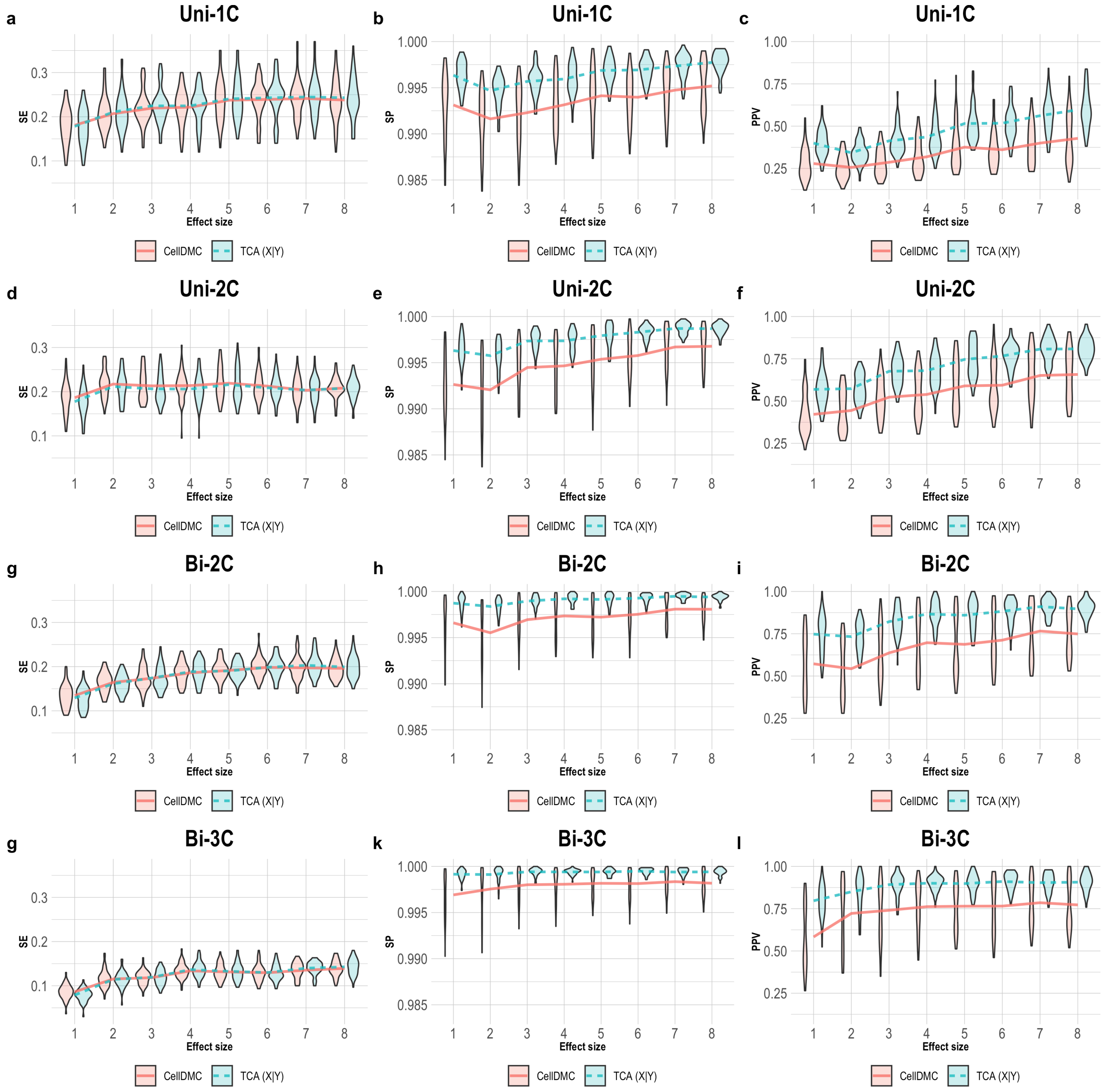

Figure 1: Evaluation of TCA and CellDMC in the case where the phenotype is affected by methylation ($Y|X$), while executing TCA under the wrong assumption $X|Y$ (TCA $X|Y$). (a)-(c) Comparison of the sensitivity (SE), specificity (SP), and precision (positive predictive value; PPV) to detect differentially methylated cell-types as a function of the association effect size, under the scenario where a single cell type out of 6 blood cell types is altered in cases versus controls (Uni-1C). (d)-(f) as in Uni-1C, only for the scenario where two cell types are altered in the same direction (Uni-2C). (g)-(i) as in Uni-2C, only for the scenario where the cell types are altered in opposite directions (Bi-2C). (j)-(l) as in Bi-2C, only for three cell types (Bi-3C). Results are shown across 50 simulated datasets using violin plots; solid lines represent median values.

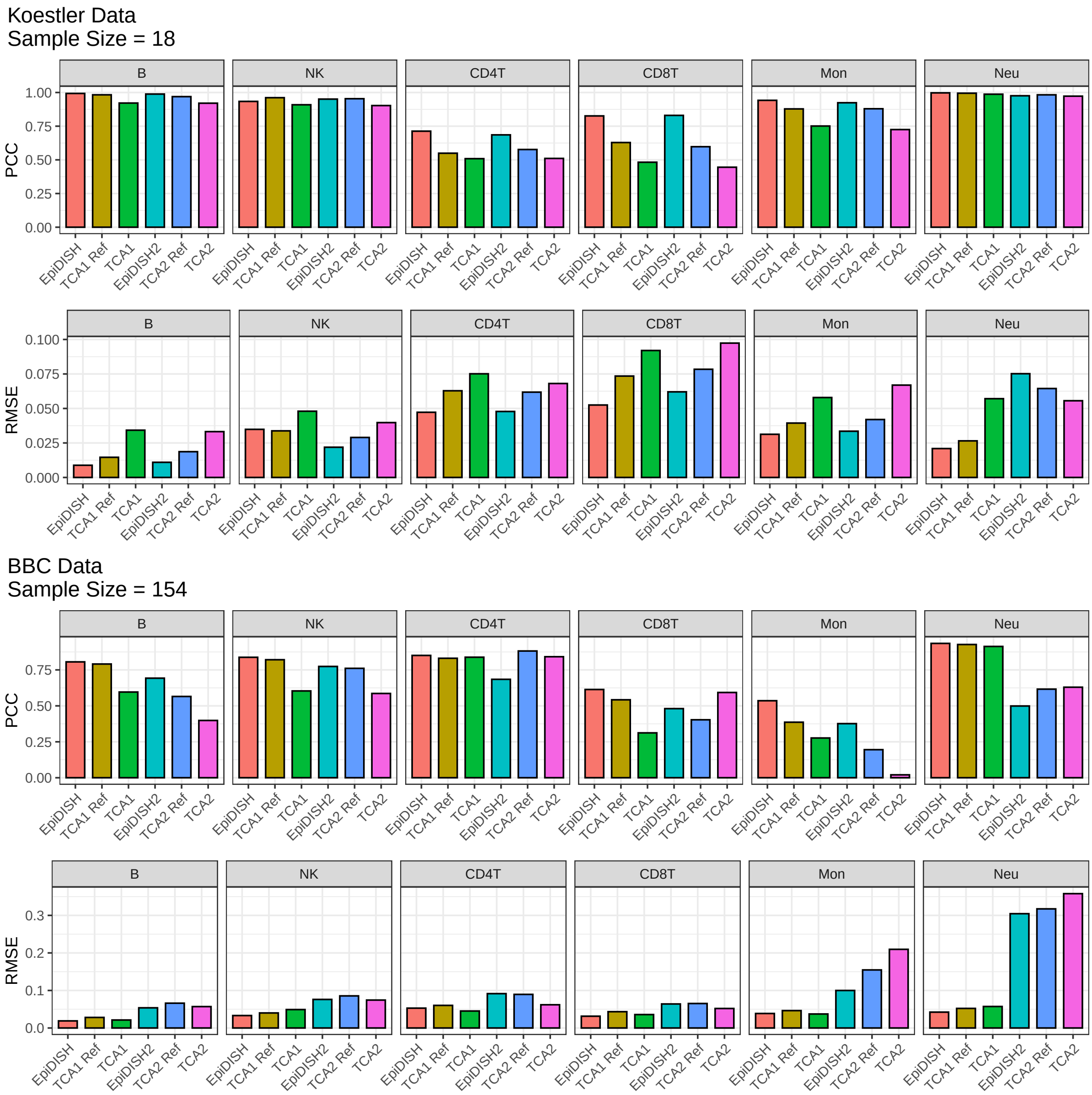

Figure 2: Evaluation of cell-type fraction estimates using TCA and a reference-based method (EpiDISH). For each of two whole-blood datasets, the data by Koestler et al. (n=18) [1] and the BBC data presented in Jing et al. (n=154), we provide barplots of the Pearson Correlation Coefficient (PCC; top panel for each dataset) and the Root Mean Square Error (RMSE; bottom panel for each dataset) between the estimated cell-type fractions and the true fractions of 6 immune cell types. PCC and RMSE are presented for three different alternatives: EpiDISH ('EpiDISH'), TCA while initializing it with EpiDISH and using the same 333 reference methylation sites used by EpiDISH ('TCA1 Ref'), TCA while initializing in with EpiDISH and using the default feature selection for model fitting ('TCA1'). In addition, these three alternatives are re-evaluated after mixing the EpiDISH estimates with noise, thus emulating the case of providing cell-type proportion estimates of lower quality as an input to TCA ('EpiDISH2', 'TCA2 Ref', and 'TCA2').

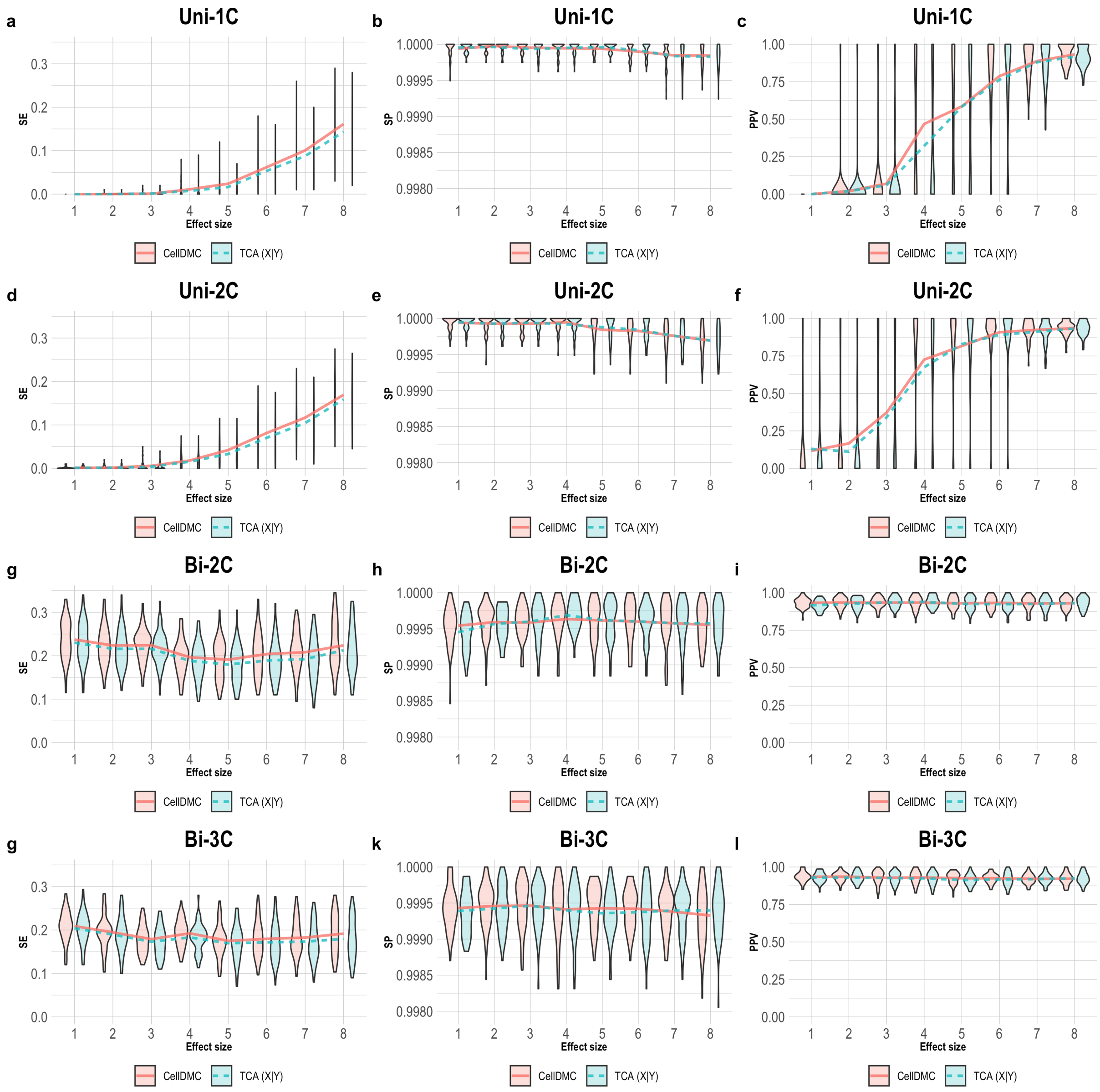

Figure 3: Evaluation of TCA and CellDMC in the case where the phenotype affects methylation $(X|Y)$ using small simulated data (n=30). (a)-(c) Comparison of the sensitivity (SE), specificity (SP), and precision (positive predictive value; PPV) to detect differentially methylated cell-types as a function of the association effect size, under the scenario where a single cell type out of 6 blood cell types is altered in cases versus controls (Uni-1C). (d)-(f) as in Uni-1C, only for the scenario where two cell types are altered in the same direction (Uni-2C). (g)-(i) as in Uni-2C, only for the scenario where the cell types are altered in opposite directions (Bi-2C). (j)-(l) as in Bi-2C, only for three cell types (Bi-3C). Results are shown across 50 simulated datasets using violin plots; solid lines represent median values. TCA was executed under the assumption $X|Y$ (TCA $X|Y$).

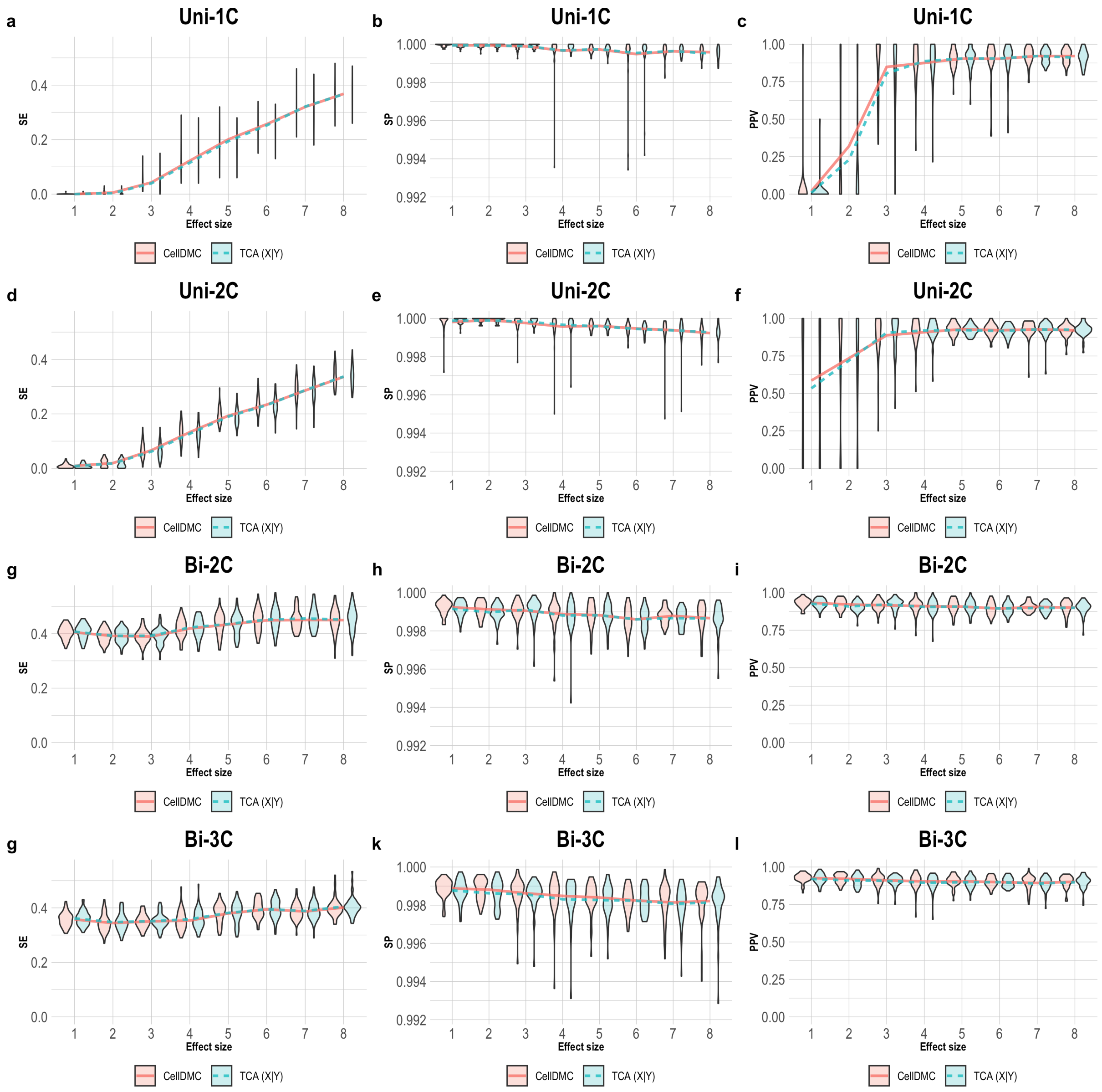

Figure 4: Evaluation of TCA and CellDMC in the case where the phenotype affects methylation ($X|Y$) using small simulated data (n=60). (a)-(c) Comparison of the sensitivity (SE), specificity (SP), and precision (positive predictive value; PPV) to detect differentially methylated cell-types as a function of the association effect size, under the scenario where a single cell type out of 6 blood cell types is altered in cases versus controls (Uni-1C). (d)-(f) as in Uni-1C, only for the scenario where two cell types are altered in the same direction (Uni-2C). (g)-(i) as in Uni-2C, only for the scenario where the cell types are altered in opposite directions (Bi-2C). (j)-(l) as in Bi-2C, only for three cell types (Bi-3C). Results are shown across 50 simulated datasets using violin plots; solid lines represent median values. TCA was executed under the assumption $X|Y$ (TCA $X|Y$).

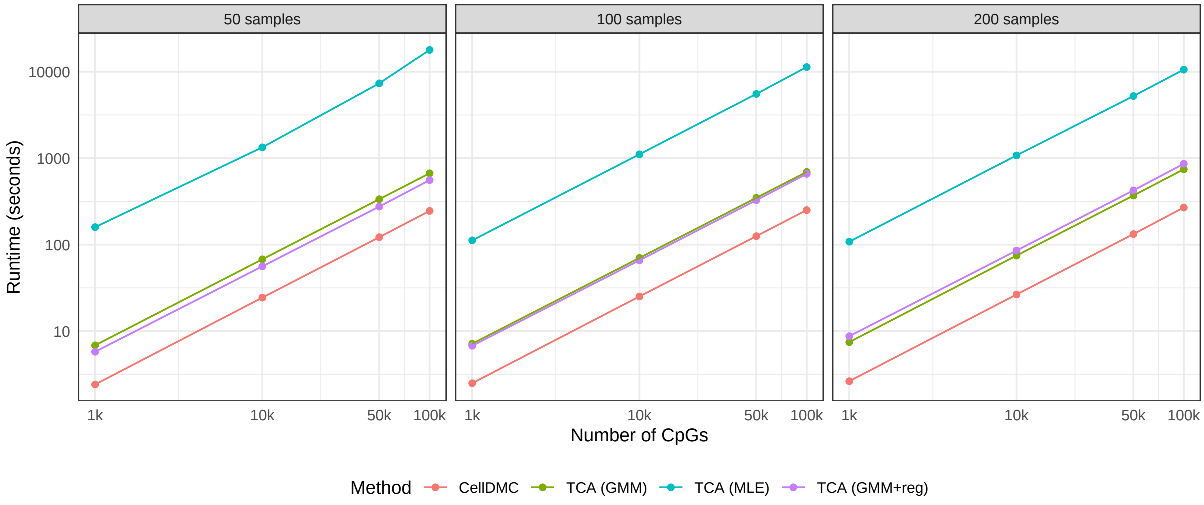

Figure 5: Evaluation of the runtimes of TCA and CellDMC. Shown are single-core runtimes (in seconds; shown in a log-base-10 scale) of CellDMC, TCA under the assumption $X|Y$ using the original approach for model fitting (using maximum likelihood; 'TCA MLE'), and TCA using the generalized method of moments as an alternative approach for model fitting ('TCA GMM' under the assumption $X|Y$ and 'TCA GMM+reg' under the assumption $Y|X$). Results are presented for each of three sample sizes (50, 100, and 200 individuals; generated using the 'test_data' function in the TCA package) as a function of the number of tested CpGs.

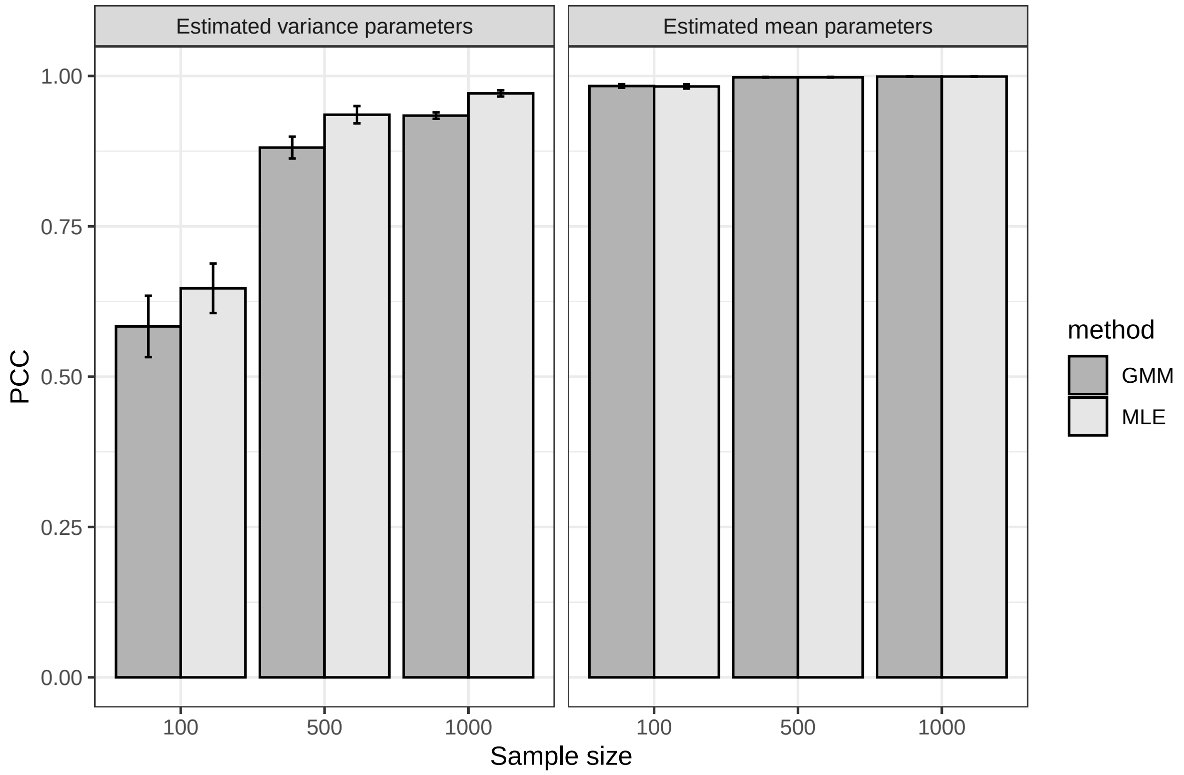

Figure 6: Evaluation of the original versus the alternative model fitting options in TCA. Barplots present the Pearson Correlation Coefficient (PCC) of true variance parameters (left) and true mean parameters (right) with their estimates as given by the original optimization of the TCA model (MLE) and the alternative optimization procedure (GMM). Results show the median performance across five simulated datasets with error bars indicating the range of performance.

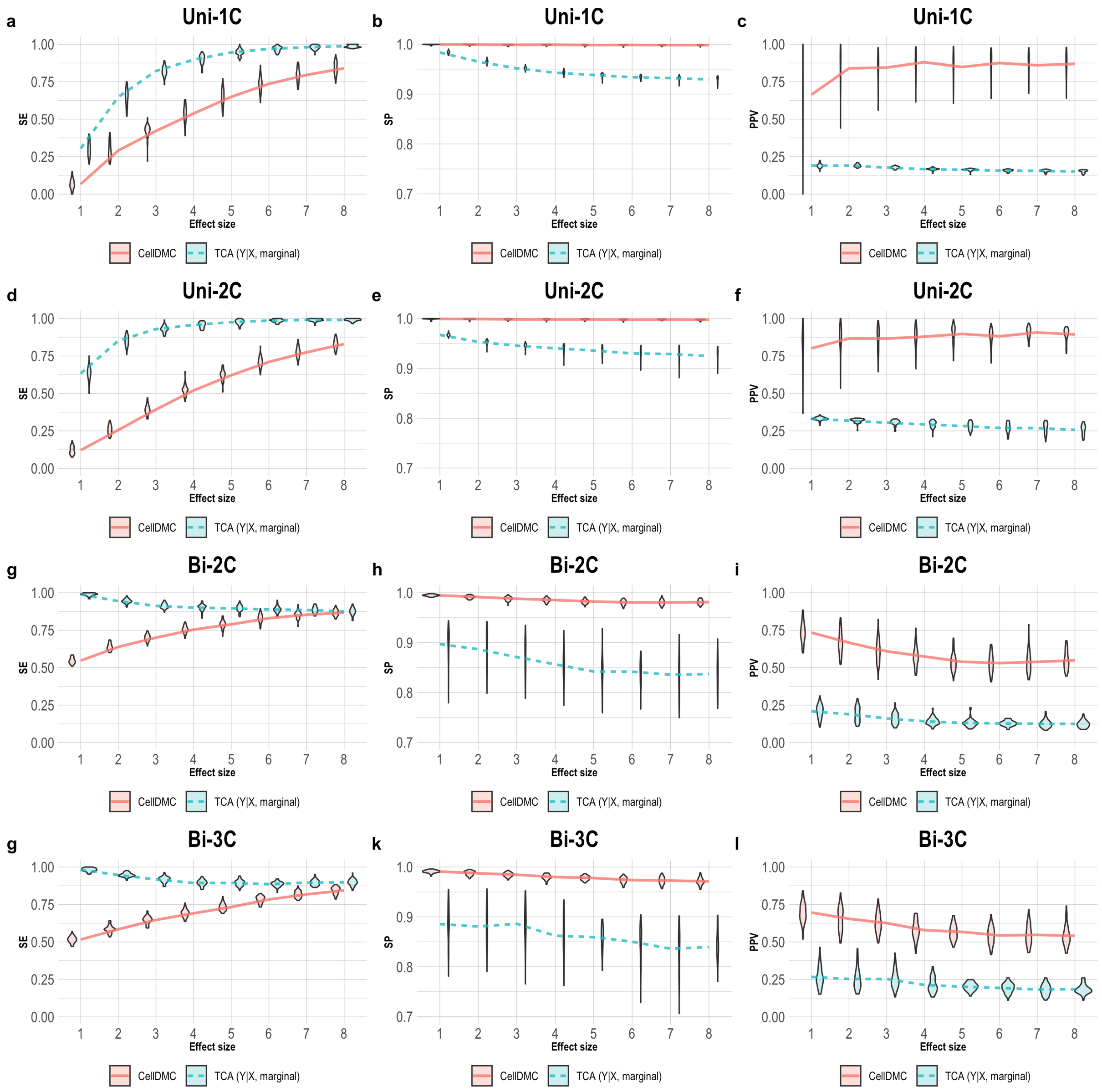

Figure 7: Evaluation of TCA and CellDMC in the case where the phenotype affects methylation $(X|Y)$, while executing TCA under the assumption $Y|X$ and using a marginal test. (a)-(c) Comparison of the sensitivity (SE), specificity (SP), and precision (positive predictive value; PPV) to detect differentially methylated cell-types as a function of the association effect size, under the scenario where a single cell type out of 6 blood cell types is altered in cases versus controls (Uni-1C). (d)-(f) as in Uni-1C, only for the scenario where two cell types are altered in the same direction (Uni-2C). (g)-(i) as in Uni-2C, only for the scenario where the cell types are altered in opposite directions (Bi-2C). (j)-(l) as in Bi-2C, only for three cell types (Bi-3C). Results are shown across 50 simulated datasets using violin plots; solid lines represent median values.

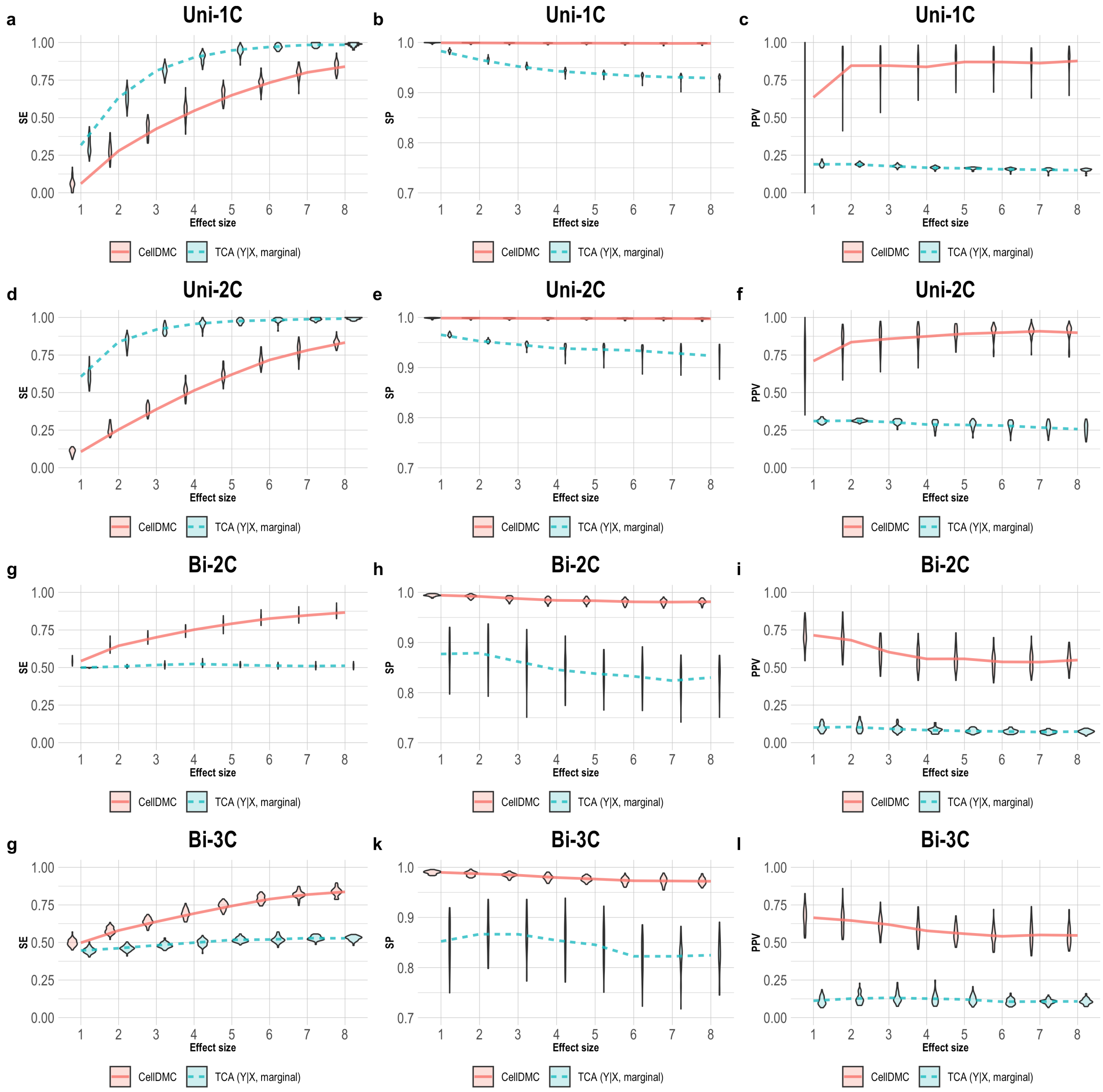

Figure 8: Evaluation of TCA and CellDMC in the case where the phenotype affects methylation $(X|Y)$, while executing TCA under the assumption $Y|X$ and using a marginal test, and while considering an additional criterion on the direction of effect in the definition of sensitivity and specificity (Supplementary Methods). (a)-(c) Comparison of the sensitivity (SE), specificity (SP), and precision (positive predictive value; PPV) to detect differentially methylated cell-types as a function of the association effect size, under the scenario where a single cell type out of 6 blood cell types is altered in cases versus controls (Uni-1C). (d)-(f) as in Uni-1C, only for the scenario where two cell types are altered in the same direction (Uni-2C). (g)-(i) as in Uni-2C, only for the scenario where the cell types are altered in opposite directions (Bi-2C). (j)-(l) as in Bi-2C, only for three cell types (Bi-3C). Results are shown across 50 simulated datasets using violin plots; solid lines represent median values.

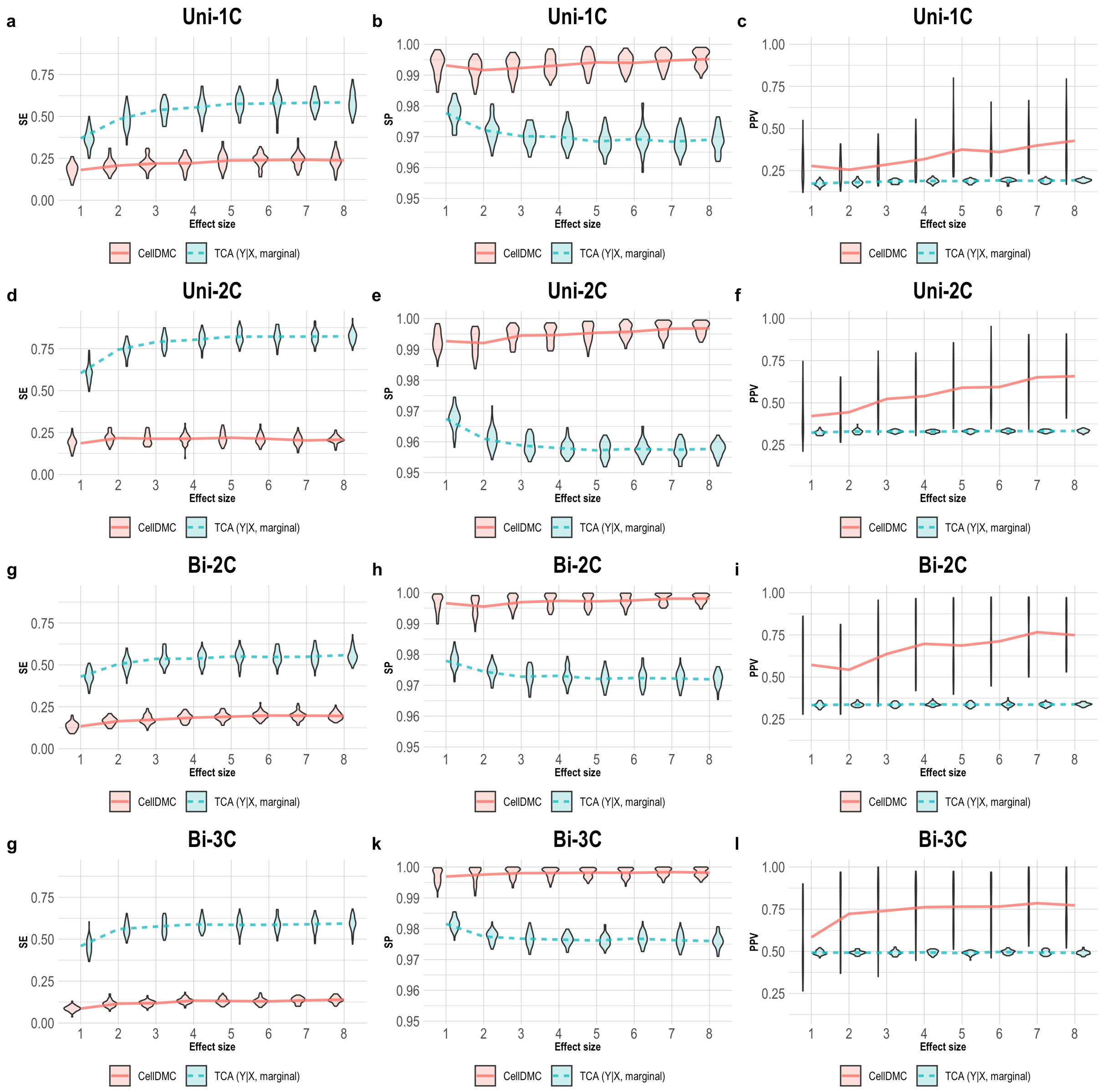

Figure 9: Evaluation of TCA and CellDMC in the case where the phenotype is affected by methylation ($Y|X$), while executing TCA under the assumption $Y|X$ and using a marginal test. (a)-(c) Comparison of the sensitivity (SE), specificity (SP), and precision (positive predictive value; PPV) to detect differentially methylated cell-types as a function of the association effect size, under the scenario where a single cell type out of 6 blood cell types is altered in cases versus controls (Uni-1C). (d)-(f) as in Uni-1C, only for the scenario where two cell types are altered in the same direction (Uni-2C). (g)-(i) as in Uni-2C, only for the scenario where the cell types are altered in opposite directions (Bi-2C). (j)-(l) as in Bi-2C, only for three cell types (Bi-3C). Results are shown across 50 simulated datasets using violin plots; solid lines represent median values.

### 3 Supplementary Note: calling differential methylation at cell-type resolution

#### 3.1 A brief overview of previous work

Genomic markers are known to demonstrate differences between different cell types. Yet, differential analysis in genomics is typically performed using tissue-level bulk data, owing to high costs and practical limitations with generating large-scale data at cell-type-specific resolution. This has led to the development of computational methods that aim at performing differential analysis experiments at the cell-type level using tissue-level data.

A first attempt in that direction suggested to estimate differential expression at a cell-type level by solving a separate decomposition problem in each of two populations of interest (e.g., cases and controls for a particular condition) [2]. A later work, which essentially further generalized this approach, allowed to consider non-categorical phenotypes by using a standard linear regression framework with interaction terms between cell-type proportions and a phenotype and interest [3]. More recently, the same interaction model was suggested and applied independently by several groups for calling differential DNA methylation at the cell-type level [4–6]. Particularly, it was suggested as a method called CellDMC by Zheng et al. [4].

A different approach that was suggested by us, called Tensor Composition Analysis (TCA), models cell-type-specific variation as individual-specific, thus assuming that the two-dimensional input data (individuals by CpGs in the case of methylation) is coming from a three-dimensional structure (individuals by CpGs by cell types). Notably, this perspective was implicitly suggested independently by another group as well at the same time [7], although, as we later discuss, they did not consider the entire span of aspects that were presented as part of the TCA framework.

Below, we go over the technical details of the decomposition approach for detecting differential methylation at the cell-type level. Then, we describe the TCA approach and relate it to the more standard decomposition approach, as well as discuss model directionality and statistical testing within the TCA framework. Finally, in the last section, we provide a practical guide with discussion about appropriate selection of model directionality and statistical tests for analysis.

##### 3.2 A two-way decomposition for differential analysis with binary phenotypes

Let $X \in \mathbb{R}^{n \times m}$ be tissue-level bulk data collected from $m$ features in $n$ individuals. A standard decomposition problem assumes the following model:

$$X = W^T Z + E \tag{1}$$

Here, $W \in \mathbb{R}^{k \times n}$ is a matrix containing for each individual their fractions of each of $k$ different cell types assumed to compose the tissue from which $X$ was collected, $Z \in \mathbb{R}^{k \times m}$ is a matrix with cell-type-specific signatures for each of the $k$ cell types in each of the $m$ features, and $E$ represents noise.

Many different versions that differ in their assumptions on the components of the decomposition (i.e. $W, Z, E$) have been proposed, and numerous successful applications in and beyond genomics employed a decomposition approach. Critically, estimating $W, Z$ jointly under the model in (1) (unsupervised decomposition) may not be identifiable, even when introducing certain constraints on the solution of the decomposition (such as non-negativity constraints and requiring the cell-type fractions of each individual to sum up to 1) [8]. To circumvent this, one can learn and fix either $Z$ or $W$ by using external reference data collected from purified cells for the former or, alternatively, measuring cell counts for the individuals in the data for the latter; this supervised decomposition approach allows to avoid the non-identifiability issue (supervised decomposition) [8, 9].

Solving a decomposition problem provides us with cell-type-specific signatures that are shared across all individuals in the data; thus, essentially assuming that all individuals have the exact same values at the cell-type level. This assumption, which is known to be unjustified biologically, is inherent in the classical decomposition formulation. Consequently, this model does not allow us to interrogate variance at the cell-type level, let alone to perform differential expression at the cell-type level. The first relaxation of this assumption (that we are aware of) in genomics was proposed by Shen-Orr et al. in the context of differential expression analysis [2]. Keeping the notations above, the authors considered the following model for tissue-level expression data:

$$x_{ij} = \sum_{h=1}^{k} w_{hi} z_{hj}^{g} + \epsilon_{ij} \tag{2}$$

$$\epsilon_{ij} \sim N(0, \sigma_g^2)$$

Here, $x_{ij}$ corresponds to the expression level of gene $j$ in individual $i$ (row $i$, column $j$ in $X$), $w_{hi}$ corresponds to the fraction of cell type $h$ in individual $i$ (row $h$, column $i$ in $W$), and $z_{hj}$ corresponds to the expression of gene $j$ in cell-type $h$ (row $h$, column $j$ in $Z$). Additionally, $g \in \{0, 1\}$ allows to represent two possible values for $Z$, one for each of two groups of individuals (e.g., cases and controls) and $\epsilon_{ij}$ is assumed to be normally distributed with a possibly different variance for each group $g$.

Shen-Orr et al. experimentally measured $W$ and solved (2), which essentially corresponds to solving a supervised version of the decomposition model in (1) twice, once for each group $g$, while fixing $W$ with the measured cell counts. Provided with estimates for $\{z_{hj}^g\}_{h,j,g}$, the authors statistically tested for differences between $z_{hj}^0$ and $z_{hj}^1$ for each $h, j$, which allowed to interrogate differential expression of gene $j$ in cell type $h$ across the two groups of individuals.

##### 3.3 Regression with interaction terms as a generalized decomposition for quantitative phenotypes

The two-way decomposition approach can in principle be extended to a multi-way decomposition, where a separate decomposition model is fitted for each possible value of a categorical phenotype of interest. However, is it not immediately clear how this approach can be extended to quantitative phenotypes. As we next show, this can be addressed under a regression framework.

Westra et al. [3] employed a regression model with interaction terms in the context of differential expression as follows. Let $y$ be an $n$-length vector with phenotypic values for the same $n$ individuals as in the data matrix $X$, keeping the previous notations, the authors considered the following model for tissue-level expression:

$$x_{ij} = \sum_{h=1}^{k} w_{hi}\mu_{hj} + \sum_{h=1}^{k} w_{hi} y_i \gamma_{hj} + \epsilon_{ij} \tag{3}$$

$$\epsilon_{ij} \sim N(0, \sigma^2)$$

where $\mu_{hj}$ is the cell-type-specific signature of gene $j$ in cell type $h$, $y_i$ is the phenotypic value of individual $i$ and $\gamma_{hj}$ is the effect size - specific to gene $j$ - of the interaction (i.e. multiplicative) term between $y$ and the proportion of cell type $h$.

For a binary phenotype $y$, it is easy to see that the model in (3) can be rephrased as a two-way decomposition model as in (2):

$$\sum_{h=1}^{k} w_{hi}\mu_{hj} + \sum_{h=1}^{k} w_{hi} y_i \gamma_{hj} + \epsilon_{ij} = \sum_{h=1}^{k} w_{hi}\tilde{z}_{hj}\mathbb{I}\{y_i = 0\} + \sum_{h=1}^{k} w_{hi} z'_{hj}\mathbb{I}\{y_i = 1\} + \epsilon_{ij} \tag{4}$$

$$\equiv \sum_{h=1}^{k} w_{hi} z^{g}_{hj} + \epsilon_{ij} \tag{5}$$

where $g$ in (5) represents the possible values for $z_{hj}$, corresponding to the two values of the phenotype $y$. Of note, the component of variation $\epsilon_{ij}$ can be treated separately for each group in $y$.

The above illustrates the motivation for performing cell-type-specific differential analysis by testing the coefficients $\{\gamma_{hj}\}$ in (3) as an evidence for different effects for different values of $y$. Clearly, rephrasing the decomposition formulation as a regression problem with interactions not only allows us to consider non-categorical phenotypes, but it also allows a straightforward model fitting and statistical analysis by leveraging the well-established tools from standard regression analysis. Particularly, covariates can be readily included in the model and tested as in standard regression:

$$x_{ij} = \sum_{h=1}^{k} w_{hi}\mu_{hj} + \sum_{h=1}^{k} w_{hi} y_i \gamma_{hj} + \sum_{d=1}^{p} c_{id}\delta_{jd} + \epsilon_{ij} \tag{6}$$

Here, $c_{id}, \delta_{jd}$ are the value of the $d$-th covariate (out of a total of $p$ covariates) of individual $i$ and its corresponding effect size, which is specific to feature $j$.

The interaction model in (6) was recently presented and applied by several groups for calling differential DNA methylation at cell-type resolution [4–6]. Most notably, it was used in CellDMC by Zheng et al. [4]. We next demonstrate TCA, a second approach that can be applied to differential analysis at cell-type resolution; later, we further relate it to the traditional interaction model taken in CellDMC.

##### 3.4 TCA: a deconvolution-based solution

While the interaction model in (6) improves upon the classical decomposition model in (1) by allowing differences in cell-type-specific profiles between individuals that are coming from different groups, it is still limited by the assumption that cell-type-specific levels are constant across individuals when conditioning on the phenotype of interest (whether it is categorical or continuous). Put differently, if we consider a case/control scenario as an illustrating example, the interaction model assumes that all cases (and similarly for controls) share the exact same cell-type-specific levels. The inter-group variation under this model stems only from fixed effects of the phenotype, while ignoring possible effects of additional factors at the cell-type level as well as intrinsic variability per individual (i.e. individual-specific unexplained biological noise).

Motivated by these limitations of previous models, we recently introduced the TCA framework, in which we model the assumption that different individuals may demonstrate differences in cell-type-specific profiles owing to multiple factors and individual-specific intrinsic variability [10]. We presented TCA in the context of DNA methylation and applied it for detecting differential methylation with a cell-type-specific resolution.

In more details, the TCA model for methylation levels at the cell-type level makes the following assumption:

$$Z_{hj}^{i} = \mu_{hj} + \sum_{d=1}^{p_1} c_{id}^{(1)} \gamma_{hd}^{j} + \epsilon_{hj}^{i} \tag{7}$$

$$\epsilon_{hj}^{i} \sim (0, \sigma_{hj}^{2})$$

where $Z_{hj}^{i}$ is a random variable that represents the level in methylation site $j$ and cell type $h$ for the $i$-th individual, $\mu_{hj}, \sigma_{hj}$ are the population-level mean value and standard deviation for site $j$ and cell type $h$, and $c_{id}^{(1)}, \gamma_{hd}^{j}$ are the value of the $d$-th covariate (out of a total of $p_1$ covariates) of individual $i$ and its corresponding effect size that is specific to methylation site $j$

and cell type $h$. Note that (7) considers $p_1$ covariates that are assumed to affect methylation at the cell-type level.

The TCA model further assumes the following model for the observed tissue-level data:

$$X_{ij} = \sum_{h=1}^{k} w_{hi} Z_{hj}^{i} + \sum_{d=1}^{p_2} c_{id}^{(2)} \delta_{jd} + \epsilon_{ij} \tag{8}$$

$$\epsilon_{ij} \sim N(0, \tau^2)$$

where $\epsilon_{ij}$ is an i.i.d. component of variation and $c_{id}^{(2)}, \delta_{jd}$ are the value of the $d$-th covariate (out of a total of $p_2$ covariates) of individual $i$ and its corresponding effect size that is specific to site $j$. Note that (8) considers $p_2$ covariates that are assumed to affect the tissue-level methylation mixtures (i.e. rather than the methylation at the cell-type level; such covariates can be, for example, batch information, that may affect the mixture regardless of the cell composition and the underlying cell-type-specific signals).

The above model reflects the assumption that the two-dimensional input data ($X$; individuals by sites) is coming from a three-dimensional underlying structure ($\{z_{hj}^{i}\}_{h,j,i}$; individuals by sites by cell types). The TCA framework allows for learning the cell-type-specific levels of an individual from their tissue-level data, thus performing a deconvolution of the observed signals in $X$ into their underlying signals; of note, we make a distinction between a deconvolution that aims at obtaining the complete three-dimensional underlying signals and a decomposition that aims at obtaining population-level properties from the data, as in (1) and (6). Performing a deconvolution in this case becomes possible by the fact that both the data points and the entries of the underlying three-dimensional tensor are treated as random variables. Consequently, we can look at the conditional distribution of the tensor given the observed data and use it for inferring the tensor values of the observed data. Particularly, the TCA estimator is defined as:

$$\hat{z}_{hj}^{i} = E[Z_{hj}^{i} | X_{ij} = x_{ij}; \Theta] \tag{9}$$

where $\Theta$ represents the parameters in (7) and in (8) - these are unknown, however, they can be estimated from the data [10]. Particularly, $W$ is assumed to be known in a typical application of TCA, however, we developed an alternative optimization procedure for re-estimating $W$ from the data in the case where only low-quality estimates of the cell-type proportions are available [10].

Notably, even though the model in (6), as presented in CellDMC, does not make a distinction between cell-type-specific covariates and mixture-level covariates, it can allow so by considering interaction terms between the cell-type proportions and the covariates that are assumed to have cell-type-specific effects. Yet, TCA further handles individual-specific intrinsic variability (i.e. the component $\epsilon_{hj}^{i}$ in (7)), which is not modeled by CellDMC.

In parallel to the introduction of TCA, another group presented essentially the same model as in (7) and in (8) [7] and applied it for calling cell-type-specific differential methylation by treating the phenotype of interest as a cell-type-specific covariate. The TCA framework, however, is more general: it allows both to treat the phenotype as a covariate (i.e. as an explaining variable of methylation) and, as we later discuss, to directly model the phenotype (i.e. as the explained variable, while considering methylation levels as explaining variables). In addition, as discussed above, TCA allows to explicitly estimate cell-type-specific methylation profiles for each individual.

##### 3.5 Relating the TCA model to the interaction model (CellDMC)

In the original TCA paper, we primarily focused on directly modeling a phenotype of interest as affected by cell-type-specific methylation, while making the assumptions in (7)-(8) for the methylation levels (see details and discussion in the next section). However, since the model in (7)-(8) can take into account covariates that affect methylation levels (or mediating components thereof) at the cell-type level, the phenotype of interest can also be treated as a cell-type-specific covaraite; we previously discussed this alternative in the original TCA paper (see "Applying TCA to epigenetic association studies" in the Methods section of the original TCA paper) [10].

Let $y$ be a phenotype of interest that may affect methylation at the cell-type level, assuming no other factors affect methylation for simplicity, the TCA model can be formulated as follows:

$$Z_{hj}^{i} = \mu_{hj} + y_i \gamma_h^j + \epsilon_{hj}^{i}, \epsilon_{hj}^{i} \sim N(0, \sigma_{hj}^2) \tag{10}$$

$$X_{ij} = \sum_{h=1}^{k} w_{hi} Z_{hj}^{i} + \epsilon_{ij}, \epsilon_{ij} \sim N(0, \tau^2) \tag{11}$$

Disregarding the component of intrinsic variation at the cell-type level (i.e. $\epsilon_{hj}^i$) by assuming $\sigma_{hj} = 0$, we get:

$$z_{hj}^i = \mu_{hj} + y_i \gamma_h^j \tag{12}$$

where $z_{hj}^i$ is now a constant value conditional on $y_i, \gamma_h^j$ (and therefore we make a distinction from the notation $Z_{hj}^i$, which represents a random variable with non-zero variance). Under this assumption, the model of $X$ can be summarized as follows:

$$\begin{aligned} x_{ij} &= \sum_{h=1}^{k} w_{hi} z_{hi} + \epsilon_{ij} && (13) \\ &= \sum_{h=1}^{k} w_{hi} \mu_{hj} + \sum_{h=1}^{k} w_{hi} y_i \gamma_h^j + \epsilon_{ij} && (14) \end{aligned}$$

where $\epsilon_{ij} \sim N(0, \tau^2)$, and where, as before, we make a distinction between $x_{ij}$ here and $X_{ij}$ in (11) (where cell-type-specific components are non-trivial random variables). This model is exactly the model in (3), which is the one used in CellDMC.

We conclude that assuming no intrinsic variability at the cell-type level (i.e. setting $\sigma_{hj} = 0$) yields TCA as equivalent to the CellDMC model, which reveals CellDMC as a degenerate case of TCA. Unlike CellDMC, the generality of TCA allows it to absorb and account for intrinsic variability at the cell-type level, and for that reason, TCA is expected to perform better than CellDMC in cases where intrinsic cell-type variation exists and equally well as CellDMC in cases where intrinsic cell-type variation does not exist or is minimal.

Importantly, the above result is specific to the case where methylation is assumed to be affected by the phenotype. As we show next, the TCA framework also allows to accommodate the assumption that the phenotype of interest if statistically affected by methylation (either directly or by a mediating component). In that case, CellDMC cannot be clearly related to TCA, as it does not allow an analogous modeling of the phenotype.

##### 3.6 Changing the model directionality: modeling the phenotype within the TCA framework

All the models we discussed so far aim at explaining methylation, and differential methylation analysis is made possible within these models by testing whether a phenotype of interest statistically affect the methylation levels under test. Essentially, this is done by including the phenotype in the model of the methylation levels. In practice, however, the directionality of the true underlying model may be different. That is, the phenotype of interest may be affected by methylation levels (or by a component that is statistically captured by methylation). In those cases, the models discussed above are statistically unjustified and empirically lead to worse performance as a result of making an incorrect assumption (Figure 2 in the main text). In what follows, we denote the assumption that methylation *affects* the phenotype by $Y|X$, and we denote the assumption that methylation is *affected* by the phenotype inversely as $X|Y$.

The TCA framework allows to accommodate the assumption that methylation affect the phenotype by directly modeling the phenotype. Particularly, it considers the following model for testing a given methylation site $j$:

$$Y_i = \sum_{h=1}^{k} Z_{hj}^{i} \beta_{hj} + e_i \tag{15}$$

$$e_i \sim N(0, \phi^2)$$

Here, $Y_i$ is the phenotypic level of individual $i$ (we make a distinction from our previously defined $y_i$ to reflect the fact that the phenotype is now a function of the random variables $\{Z_{hj}^{i}\}_{h,j}$) and $\beta_{hj}$ is the effect size of the methylation level in cell type $h$ at site $j$ on the phenotype.

Notably, depending on the context and phenotype of interest, assuming $Y|X$ as in (15) may be more appropriate (and interesting) than the alternative assumption $X|Y$. As an example, consider our previous analysis from the original TCA paper, where we applied TCA to previously studied data with rheumatoid arthritis [11]. In this analysis, three of the associated CpGs we found are highly heritable (cg13081526, cg18816397, cg13778567; more than 50% of their variances can be explained by their cis-SNPs [12]). These findings are therefore consistent with the possibility that methylation mediates causal genetic effects on rheumatoid arthritis, which rationalizes the assumption $Y|X$.

Fitting the model in (15) within the TCA framework can be done in one of two ways. First, cell-type-specific methylation can be inferred following (9), which then allows to employ a standard regression analysis. Second, we can consider the conditional distribution of the phenotype given $\{Z^i_{hj}\}_{h,j,i}$, however, since these are random variables with unobserved values they need to be integrated over. We can do so by using the conditional distribution $Z^i_{hj}|X_{ij}$ as follows:

$$\int_{z_1} ... \int_{z_k} Pr\left(Y_i = y_i|\{Z^i_{hj} = z_h\}^k_{h=1}\right) Pr\left(\{Z^i_{hj} = z_h\}^k_{h=1}|X_{ij} = x_{ij}\right) dz_1...dz_k \quad (16)$$

$$= \quad Pr\left(Y_i = y_i|X_{ij} = x_{ij}\right) \quad (17)$$

Therefore, in this second approach, TCA fits the conditional model $Y_i|X_{ij} = x_{ij}$, which can be expressed in terms of the effect sizes $\{\beta_{hj}\}_{h,j}$, thus allowing to estimate and statistically test them [10].

Of note, taking either approach to fit the model in (15) still requires to obtain estimates for the parameters of the methylation model in (7)-(8) [10]. These can then be used in the first approach for estimating $\{Z^i_{hj}\}_{h,j,i}$ following (9) or in the second approach for fitting the distribution $Y_i|X_{ij} = x_{ij}$. Regardless of which approach is taken, under the $X|Y$ assumption, the phenotype should not be considered as a cell-type-specific covariate as before in the $X|Y$ assumption.

##### 3.7 Statistical testing within the TCA framework

The TCA framework allows us to run several different types of statistical tests on a phenotype of interest, each of which can test a different hypothesis about the statistical relation of the phenotype to the methylation levels under test. In this section, we briefly describe the statistical tests we implemented as part of the TCA R package (`TCA` on CRAN; Supplementary Methods). Then, in the next section, we provide a practical guide with recommendations for the selection of appropriate tests.

Much like in standard regression analysis, where we can test different hypotheses about the coefficients of the independent variables, both the model in (7)-(8) and in (15) can consider and test several different hypotheses about the effect sizes $\beta_{1j}, ..., \beta_{kj}$. Below, we provide a list of the tests we implemented in the TCA package for testing the statistical association of a given phenotype with each particular methylation site $j$ in the data.

Tests under the biological assumption $Y|X$, using the model in (15):

- *marginal conditional* - fits the parameters $\beta_{1j}, ..., \beta_{kj}$ for all cell types jointly and tests for the significance of the effect of each cell type separately.
- *marginal* - for each cell type $h$, fits the parameter $\beta_{hj}$ and tests for the significance of its effect, while assuming $\forall l \neq h : \beta_{lj} = 0$; repeats for each cell type separately.
- *joint* - fits the parameters $\beta_{1j}, ..., \beta_{kj}$ for all cell types jointly and tests for the significance of the overall effect across all cell types (i.e. a tissue-level test).
- *single effect size* - fits the parameters $\beta_{1j}, ..., \beta_{kj}$ under the assumption $\beta_{1j} = ... = \beta_{kj}$ and tests for the significance of the overall effect across all cell types.
- *custom* - compares and tests any two nested models, each representing a subset of the parameters $\beta_{1j}, ..., \beta_{kj}$, and tests for the significance of the overall effect across all cell types in the alternative model (i.e. the larger model) that are not in the null model (i.e. the smaller model).

Tests under the biological assumption $X|Y$, using the model in (7)-(8):

- *marginal conditional* - same as the analogous model for $X|Y$ above, only under the assumption $X|Y$.
- *joint* - same as the analogous model for $X|Y$ above, only under the assumption $X|Y$.

Jing et al. [13] benchmarked TCA and CellDMC under several scenarios, in all of which they applied TCA using a *marginal* test under the biological assumption $Y|X$. CellDMC, on the other hand, makes the biological assumption $X|Y$ and applies a *marginal conditional* test. These differences, in conjunction with the fact that Jing et al. used $X|Y$ as the true model in their simulations, provided a biased perspective of the performance of the two methods.

Of note, CellDMC can in principle define more statistical tests on the cell-type-specific effect sizes (i.e. note just the marginal conditional test), similarly to the tests allowed in TCA.

However, these are correctly not implemented, and even if they were implemented, we would expect these tests to be inferior to their analogous ones in TCA given that CellDMC is a degenerate case of TCA (see subsection 3.5). Additionally, there is no clear way to accommodate the assumption $Y|X$ in CellDMC, as the method relies on the interaction between cell-type proportions and a phenotype, which is the variable to be modeled under $Y|X$.

We further provide in Supplementary Table 1 a summary of the statistical tests available in both TCA and CellDMC, including the functions for running them. For more in-depth details about the statistical tests available in the TCA package refer to the package documentation and the original TCA paper [10].

##### 3.8 A practical guide: selecting appropriate statistical tests for differential methylation at cell-type resolution

The first important decision one should make when testing for differential methylation at cell-type resolution is how to set model directionality. While this decision clearly should be context- and phenotype-dependent, admittedly, it may often not be clear how to do make an informed decision. Yet, in some cases, the selection of a biological assumption is natural. For example, when looking for associations with demographic factors such as age or ancestry, it makes no sense to assume $Y|X$, as these demographics cannot be altered by methylation.

Based on the results from our simulation study, the consistency between TCA and CellDMC may provide further useful evidence as for the true underlying model. Specifically, high consistency in the predicted associations between TCA and CellDMC while applying TCA under the assumption $X|Y$ provides evidence that the assumption $X|Y$ holds (Figure 1 in the main text). In contrast, limited consistency between the two methods owing to lower specificity and precision of CellDMC - which is expected to results in more predicted associations for CellDMC over TCA - provides evidence that the assumption $Y|X$ holds; either when applying TCA under the assumption $X|Y$ (Supplementary Figure 1) or when applying TCA under the assumption $Y|X$, which is expected to provide even worse inconsistency based on our simulations (Figure 2 in the main text). That said, in practice, the extent to which a given phenotype will demonstrate patterns of consistency that are similar to those revealed by simulations is still unclear; particularly, a phenotype may be both affected by some methylation sites (or by a component captured by methylation) and affect some other methylation sites (or a mediating component thereof).

In cases of association studies that aim at de-novo detection of differential methylation, we recommend to apply a joint test for an initial screening for tissue-level associations. Then, marginal conditional tests should be performed as a post-hoc analysis for calling differentially methylated cell types in the CpGs that passed multiple testing correction in the first tissue-level screening. As per our previous suggestion, we recommend that future studies include small replication data sets from sorted or single cells, in which case users may opt to replace marginal conditional tests with the much more powerful, yet less precise marginal tests; in such cases, pre-screening for tissue-level associations using joint tests may be less powerful.

In cases where only a single cell type (or a small subset of cell types) is associated with the phenotype, joint tests are expected to be less powerful than marginal and marginal conditional tests (owing to the unnecessarily higher degrees of freedom in a joint test). While this is typically unknown a priori, this rationale can be applied to cases where only a particular cell type (or a small subset of cell types) are of interest, in which case, an initial screening step using a joint test should be avoided.

Lastly, it is important to understand the limitations of methods such as TCA and CellDMC. Particularly, there are limitations that rise due to inherent properties of these methods: first, the proportions of different cell types are correlated, owing to the fact that fractions sum up to 1 and thus depend on each other, and second, the higher the abundance of a cell type is, the higher the variance that it accounts for in the observed mixture data. Consequently, the estimated cell-type-specific methylation in TCA and the cell-type-phenotype interactions in CellDMC, both of which directly rely on the cell-type proportions, are expected to be correlated between different cell types. For that reason, these two models are expected to be bounded in their precision to detect truly differentially methylated cell types.

In order to see that, consider an example where we have cell type A with accurate estimates of the cell-type proportions and cell type B with less accurate estimates of the cell-type proportions (e.g., granulocytes and monocytes in whole-blood). Further assume that cell type B is truly differentially methylated at some particular CpG under test. In that case, if the proportions of both cell types are highly correlated (and they typically are), cell type A may capture some of the cell-type-specific methylation of cell type B that was not captured by directly using the proportions of cell type B (due to the limited accuracy of the estimated proportions of cell type B and the correlation between the proportions of A and B). Not only that, these correlations between the proportions of different cell types may introduce high multicollinearity between estimated methylation of different cell types and their effects. As a result of these, cell type A in our example may be called as differentially methylated, even though the true signal is coming from cell type B.

The above limitations are also the reason for the particularly low precision of the marginal test in TCA, where the cell type under test tends to capture true signal coming from other cell types as well (Figures 7, 8, and 9), much like what one would observe when applying marginal tests under standard linear regression in the case of having multiple highly correlated features (where only some of them are truly statistically related to the dependent variable). Importantly, applying a marginal conditional test instead of a marginal test mitigates this limitation (although not completely, as explained above) by accounting for the other cell types.

Finally, we note that the above inherent limitation of the marginal test motivates the exclusion of the requirement that estimated effect sizes of predicted associations match true effects when evaluating the sensitivity of the marginal test under simulations (a criterion which was used by Jing et al. [13]), assuming the question of interest is solely the detection of differentially methylated cell types, regardless of the direction of effects.

### 4 Supplementary Methods

#### 4.1 Software and computational tools

We applied TCA using the `TCA` R package version 1.2.0 (available on CRAN); source code is available from github at github.com/cozygene/TCA. For the application of CellDMC, we used the `CellDMC` function in the EpiDISH R package version 2.2.0 (available on Bioconductor) which provides an implementation of CellDMC.

In the application of TCA under the assumption $X|Y$, we used the `tca` function, which now provides p-values for the estimated parameters in the model under a marginal conditional test and under a joint test; these are given in the output fields `gammas_hat_pvals` and `gammas_hat_pvals.joint` of the `tca` function. In the application of TCA under the assumption $Y|X$, we used the `tcareg` function, while setting the argument `test` to the requested type of test (e.g., marginal or marginal conditional). Throughout our experiments, unless stated otherwise, we used the fast mode of the `tca` function by setting `vars.mle = FALSE` and the fast mode of the `tcareg` function by setting `fast_mode = TRUE` (see next subsection for details).

Finally, we provide an R package called `CellTypeSpecificMethylationAnalysis`, which allows a complete reconstruction of our experiments and figures. The full source code of the package is available from github at: github.com/cozygene/CellTypeSpecificMethylationAnalysis.

#### 4.2 Fast optimization of the TCA model

We previously applied an alternating maximum-likelihood-based optimization procedure for fitting the TCA model [10]. Learning the mean parameters in the model in equations (7)-(8) given the variance parameters is a convex problem that can be solved efficiently by formulating it as a constrained regression problem, yet, estimating the variance parameters (i.e. given estimates of the means) is a non-convex problem. For that reason, we employed a gradient-based optimization for the variances. This resulted in relatively long runtimes of the function `tca` in the TCA R package, especially for large data such as methylation arrays that include hundreds of thousands of features.

Maximum-likelihood estimation is perhaps the most common approach for fitting statistical models, however, alternatives do exist. Particularly, the generalized method of moments (GMM) allows to estimate model parameters by defining moment conditions, which are essentially sets of equations that are constructed from the model parameters and the data [14]. Given that several assumptions on the moment conditions are met, parameter estimation with proven statistical properties can then be performed by solving efficient quadratic programming problems [14,15].

We applied the GMM technique for a fast estimation of the variances in the TCA model. Particularly, for each methylation site, we estimated the cell-type-specific variance parameters of all cell-types jointly. In order to do so, for each site, we define a set of moment conditions, one per each individual sample in the data. Each such individual-based moment condition formulates an estimator for the variance of the methylation level of the individual in the particular site under consideration. Since the variance of each given individual is a function of both the individual-specific cell-type proportions and the variance parameters of all cell types, these moment conditions can be used to estimate the variance parameters under the GMM framework [14].

We updated the `tca` function in the TCA package to include an argument `vars.mle`, which can set `tca` to use either the original maximum-likelihood estimation procedure for the variances (if set to `TRUE`) or the alternative, GMM-based procedure (if set to `FALSE`). We further verified that both alternatives perform similarly in estimating the model parameters (Figure 6). Importantly, setting `vars.mle = FALSE` allows a dramatic reduction in runtime (by almost two orders of magnitude), which results in a computation cost that is almost in par with the efficient regression model suggested in CellDMC (Figure 5).This allows us to efficiently conduct large-scale epigenome-wide studies using the `tca` function, that is, under the assumption that methylation is affected by the phenotype of interest (i.e. $X|Y$; see subsection 3.4).

The TCA framework further allows for statistical testing under the assumption that cell-type-specific methylation affect the phenotype of interest or a mediating component thereof (i.e.

$Y|X$; see subsection 3.6); this assumption is implemented in the TCA package within the `tcareg` function. As discussed in the original TCA paper, there are two approaches to perform statistical testing under the assumption $Y|X$. First, we can use a two-step approach of obtaining estimates for the cell-type-specific levels (using the `tensor` function in the TCA package), which can then be associated with the phenotype under a standard regression framework. Second, we can consider a single-step approach of directly using the conditional distribution of the phenotype given the data for statistical inference and testing (see subsection 3.6).

Previously, we implemented only the one-step approach in `tcareg`. In order to streamline the faster statistical testing that is allowed by the two-step approach, we updated the `tcareg` function accordingly to include an argument `fast_mode`, which can set `tcareg` to use either the previously implemented one-step approach (if set to `FALSE`) or the faster two-step approach (if set to `TRUE`).

##### 4.3 Simulation study

We designed our simulations based on the simulation setup suggested by Jing et al. [13] as follows. For each dataset we simulated, we generated tissue-level bulk data for a subset of the methylation sites that are available in the Reinius et al. data [16]: 1,000 sites picked at random and an additional set of 333 reference CpGs that are used in the software EpiDISH for reference-based estimation of cell-type proportions [17]. For simulating methylation levels of an individual, we first sampled cell-type-specific methylation levels for the 1,333 sites in each of six major immune cell types (CD4+, CD8+, granulocytes, monocytes, B cells, and natural killer cells) using Beta distributions that we learned from the purified methylation profiles of these cell types in the Reinius et al. data (n=6 for each cell type) [16]. Eventually, we constructed tissue-level methylation values by linearly mixing them according to cell-type proportions that we sampled from a pool of estimates we obtained by applying the reference-based method EpiDISH to the Hannum et al. data [18].

In the experiments under the assumption $X|Y$ (i.e. methylation is affected by the phenotype), unless otherwise stated, each dataset we generated was consisted of 500 individuals (to reflect a typical sample size in association studies), out of which 250 were cases and 250 controls. For simulating differentially methylated cell-types, we first selected 100 sites and cell types at random (the number of cell types was determined by the specific scenario under consideration as explained later), while requiring the selected sites to exhibit either low (<0.2) or high (>0.8) average methylation levels in the specific cell-types to be altered (the former were used for simulating hypermethylation in cases and the latter for hypomethylation in cases). Then, we altered the cell-type-specific methylation of cases in the selected sites and cell types based on the following equations, per the suggestion of the authors in Jing et al.:

$$\gamma = \frac{|\mu_1 - \mu_2|}{\sqrt{\frac{\sigma_1^2+\sigma_2^2}{2}}} \tag{18}$$

$$\sigma_1 = \sqrt{\frac{\mu_1(1-\mu_1)}{\mu_2(1-\mu_2)}}\sigma_2 \tag{19}$$

Here, considering one particular differentially methylated cell type in a given methylation site, $\gamma$ denotes the effect size and $\{\mu_1, \sigma_1^2\}, \{\mu_2, \sigma_2^2\}$ denote sets of the mean and variance of the methylation levels of the particular site and cell type in the cases and controls groups, respectively. Setting $\mu_1, \sigma_1$ was done using the above equations given a particular effect size under consideration and given the parameters $\{\mu_2, \sigma_2\}$, which were set to the mean and variance of the beta distributions that were estimated based on the Reinius et al. data as explained above. Given $\{\mu_1, \sigma_1^2\}$, these were considered as the mean and variance of a beta distribution from which methylation levels were sampled for cases.

In the experiments under the assumption $Y|X$ (i.e. methylation affects the phenotype), unless otherwise stated, each dataset we generated was consisted of 500 individuals; these were similarly generated as in the $X|Y$ simulations, with the exception that methylation levels of a specific cell type in a specific site were sampled from a single beta distribution (i.e. the same distribution for all individuals, as opposed to the separate distribution for cases and controls in the $X|Y$ simulations). We selected differentially methylated sites and cell types as performed previously in the $X|Y$ simulations, and then simulated a phenotype for each differentially methylated site

$j$ as follows:

$$y_i^j = w_i^T \alpha_j + \sum_{h \in S_j} Z_{hj}^i \beta_{hj} + \epsilon_{ij} \tag{20}$$

$$\alpha_j \sim N(0, 1) \tag{21}$$

$$\epsilon_{ij} \sim N(0, \sigma_j^2) \tag{22}$$

$$\sigma_j^2 = \sum_{h \in S_j} \sigma_{hj}^2 \tag{23}$$

where $y_i^j$ is the phenotypic value of individual $i$, $w_i$,$\alpha_j$ are the individual's cell-type proportions and their corresponding effect sizes in site $j$, respectively, $S_j$ is the set of all differentially methylated cell types in site $j$, $Z_{hj}^i, \beta_{hj}$ are the cell-type-specific methylation of individual $i$ in site $j$ and cell type $h$ and the corresponding effect size, respectively, and $\sigma_{hj}$ is the standard deviation of cell type $h$ in site $j$.

The rest of the simulated sites that are not differentially methylated were tested against one of the 100 simulated phenotypes (selected at random). Notably, equation (23) allows a meaningful definition of effect sizes by accounting for changes in variability between different methylation sites and cell types. Further, $\beta_{hj}$ can be either positive or negative under the above formulation, however, in our final evaluations (i.e. in the figures) we consider the absolute value of $\beta_{hj}$ as the effect size.

In both the $X|Y$ and $Y|X$ experiments, we considered all four scenarios that were suggested by Jing et al.: unidirectional change in one cell type, unidirectional changes in two cell types, bidirectional changes in two cell types, and bidirectional changes in three cell types. For evaluation metrics, we considered specificity, and precision (positive predictive value ; PPV) as suggested by Jing et al. We further considered sensitivity as in Jing et al., however, with a single exception. Jing et al. required that the direction of a predicted association must match the direction of the true effect in order to be counted as a true positive (and otherwise they considered it as a false positive). We disregarded this criterion for reasons that are related to the inherent multicolinearity in TCA and CellDMC (see subsection 3.8). Yet, we did rerun all experiments while taking this additional criterion into account in the definition of sensitivity, and we found that results are essentially the same in all experiments (and conclusions therefore remain the same throughout the paper; data not shown), with the exception of marginal tests in the bi-directional scenarios, wherein the performance of sensitivity is greatly affected by the inclusion of this criterion (Figures (7) and (8)).

##### 4.4 Analysis of smoking status

We reanalyzed the data presented in Jing et al. [13] as follows. We obtained the normalized Illumina 450k data by Liu et al. (n=689) [11] and by Hannum et al. (n=656) [18] from the Gene Expression Omnibus (GEO; accession numbers GSE42861 and GSE40279, respectively); smoking information is not available on the GEO record of the Hannum et al. data and can be obtained from the authors. We removed two samples with no smoking information from the Liu et al. data and 66 sample with no smoking information from the Hannum et al. data, and in both datasets we defined the smoking status as a categorical variable with three categories: never-smokers, ex-smokers, and current smokers (occasional smokers were considered as smokers).

In each of the two datasets, we tested the smoking status for association with each of the seven cell-type-specific associations that were reported by Su et al. [19], which were considered in Jing et al. as the ground truth for evaluation. In our analysis, we accounted for rheumatoid arthritis status, gender, and age in the Liu et al. data, and for age, gender, and plate in the Hannum et al. data. In addition, in order to account for technical variation in the data, we considered a previously suggested approach by Lehne et al. for capturing technical variation in the Illumina 450k methylation arrays [20], wherein principal components (PCs) are calculated from control probes that are not expected to exhibit any biological signal. Specifically, in our case, we included in the analysis of each dataset the top ten PCs calculated from a set of 1,000 sites that demonstrate the lowest variance in the data (and are therefore expected to exhibit no true biological variation). Finally, for evaluating genome-wide calibration, we tested all the methylation sites in the data for association with the smoking status, except for polymorphic probes, non-specific probes, and probes of sites that are on non-autosomal chromosomes as previously suggested [21].

Our evaluation included three methods: CellDMC, TCA under the assumption $X|Y$ using marginal conditional tests, and TCA under the assumption $X|Y$ using joint tests (as opposed to the execution of TCA by Jing et al. under the assumption $Y|X$ and using marginal tests). All methods were executed under the assumption of two cell-types (myeloids and lymphoids) as in Jing et al. To that end, cell-type proportions of seven blood cell types were estimated using the reference-based method EpiDISH [17], and then aggregated within each of the lymphoid and myeloid compartments.

##### 4.5 Estimating cell-type proportions under the TCA model

We used two datasets with matched FACS cell counts: the Illumina 450k dataset by Koestler et al. (n=18; GEO accession number GSE77797) [1], and the EPIC array data presented in Jing et al. (n=162) [13]. We processed the EPIC array data using the ENmix R package [22]; specifically, we normalized the data using the `preprocessENmix` and `norm.quantile` functions and corrected for probe design type bias correction using the function `rcp`. Removing samples with any missing cell count values, as well as two observations with missing gender information, resulted in a total of 156 samples left with complete FACS counts.

We considered three different approaches for estimating cell-type proportion: (1) the reference-based method EpiDISH [17], TCA while including the EpiDISH estimates as an input (using the function `tca` while setting `refit_W=TRUE`) and informing TCA to consider only the same 333 reference CpGs that are used in EpiDISH (i.e. informing TCA with the identity of the reference sites, without the reference methylation; using the argument `refit_W.features` in the `tca` function), and (3) the same as (2) with the exception that the set of CpGs to be used was selected in an unsupervised manner using the algorithm ReFACTor [23]. The latter is the default feature selection in `tca` when setting it to re-estimate cell-type proportions (i.e. setting `refit_W=TRUE`). We further reevaluated all approaches above under the scenario where the input estimates given to TCA are noisy. To that end, we added random noise to the EpiDISH estimates and repeated the analysis using, as before, both TCA with the 333 reference CpGs and TCA using ReFACTor for feature selection.

In our application of TCA to the EPIC array data, we considered known batch information as well as ten PCs that capture technical variation as covariates (as in subsection 4.4; these were given as the argument `C2` to the function `tca`). In the case of the Koestler data (n=18), we only considered a single covariate of known batch information, as multiple covariates should not be included in very small data. In our application of ReFACTor for feature selection we similarly accounted for the same factors, and, in addition, we excluded polymorphic probes, non-specific probes, and probes of sites that are on non-autosomal chromosomes per the recommendations for the application of ReFACTor [23, 24].

As a final note, in our execution of TCA in these experiments we did not consider covariates that may affect methylation at the cell-type level (i.e. argument `C1` in the `tca` function). In general, some biological covariates, such as age and gender, may affect methylation at the cell-type level and should therefore be included in the analysis. However, considering each such covariate in the TCA model results in $k$ additional parameters (per each site; $k$ being the number of cell types). Therefore, for relatively small datasets as the ones analyzed in these experiments, we refrain from considering cell-type-specific covariates that may lead to model overfitting and, as a result, worse performance
